# Propagation of neuronal micronuclei regulates microglial states

**DOI:** 10.1101/2023.07.31.551211

**Authors:** Sarasa Yano, Natsu Asami, Yusuke Kishi, Hikari Kubotani, Yuki Hattori, Ayako Kitazawa, Kanehiro Hayashi, Ken-ichiro Kubo, Mai Saeki, Chihiro Maeda, Kaito Akiyama, Tomomi Okajima-Takahashi, Ban Sato, Yukiko Gotoh, Kazunori Nakajima, Takeshi Ichinohe, Takeshi Nagata, Tomoki Chiba, Fuminori Tsuruta

## Abstract

Microglia, resident immune cells in the central nervous system, undergo morphological and functional changes in response to signals from the local environment and mature into various homeostatic states. However, niche signals underlying microglial development and maturation remain largely unknown. In this study, we show that neuronal micronuclei propagate microglia, followed by changing microglial states during the postnatal period. We discovered that neurons passing through a dense region of the developing neocortex give rise to micronuclei and release them into the extracellular space. Moreover, neuronal micronuclei were incorporated into microglia and affected morphological changes. Loss of the *cGAS* gene alleviates effects on micronucleus-dependent morphological changes. Notably, neuronal micronuclei-harboring microglia exhibit unique transcriptome signatures. These results demonstrate that neuronal micronuclei serve as niche signals that produce novel microglial states. Our findings provide a potential mechanism for regulating the microglial state in the early-postnatal neocortex.

## Introduction

Microglia, the resident immune cells in the central nervous system (CNS), represent the first-line immune defense in the CNS by surveillance with phagocytic scavenging and recruitment of peripheral immune cells^1–3^. In addition, a series of recent studies have established that microglia act as critical players in the development, homeostasis, aging, and a wide range of diseases in the CNS^4, 5^. During development, microglia regulate the survival and differentiation of neural progenitor cells (NPCs), migration and maturation of immature neurons, and synapse formation and elimination^6–11^. Microglia also modulate neural activity and circuits in the mature brain^12, 13^. As expected by the pivotal and wide-ranged functions, microglial abnormality has been implicated in the majority of neurodevelopmental and neurodegenerative disorders^5, 14^.

The resident immune cells in the brain consist of a diverse cell population, such as microglia, myeloid cells, and border-associated macrophages (BAMs)/CNS-associated macrophages (CAMs) (hereafter referred to as BAMs). Recent fate mapping analyses have suggested that primitive macrophages arise from erythro-myeloid progenitors (EMPs), migrate into the brain rudiment via circulatory systems at embryonic day (E) 9.5^15^, and colonize the brain. Some CD206^-^ progenitor cells colonize the brain and differentiate into microglia^16^. In addition, CD206^+^ macrophages in the ventricle infiltrate into pallium at E12.5 and convert into CD206^-^ microglia, which colonize in the cerebral cortex^17^. After colonization, microglia change their states and exhibit unique, context-dependent, gene expression profiles. Indeed, these profiles are involved in physiological and pathological conditions and affect the acquisition of various microglial functions^18^. Furthermore, BAMs, which reside in the meninges, perivascular space, and choroid plexus, also exhibit unique properties^19, 20^. Therefore, understanding the mechanism that determines microglial state is key for elucidating the brain development. Microglial states are thought to be determined by both intrinsic programs and extrinsic signals, such as cytokines, extracellular matrix, and cell-cell interactions. A remarkable aspect of microglia is their dynamic behavior over time. Usually, microglia constantly extend and retract fine-ramified processes and survey the local environments to coordinate with brain homeostasis and neural circuits^21–23^. Moreover, extracellular signals and cell-cell communication locally modulate microglial differentiation, proliferation, and homeostasis^24, 25^. Hence, it is likely that context-dependent signals, which are still largely unknown, regulate the microglial characteristics and states.

Micronuclei are small nuclei produced by a chromosome segregation errors and physical stress during cellular migration^26–28^. Micronuclei (MNs) frequently appear in cancer and are used as biomarkers^29, 30^. MNs are also associated with chromothripsis and epigenetic dysregulation^26, 31–34^. Recent studies have revealed that MNs activate the cyclic GMP-AMP synthase-stimulator of interferon genes (cGAS-STING) pathway involved in the innate immune response^35–38^. After damaging the nuclear envelope, micronuclear chromatins are released into the cytoplasm and promote tumor metastasis by activating the cGAS-STING pathway. However, besides these pathological contexts, the physiological roles of MNs have not been fully understood. Moreover, to our knowledge, the roles of MNs in cell-to-cell communications have not been described in any context.

In this study, we report that MNs are generated in migrating neurons in the developing mouse neocortex under physiological conditions. The number of micronuclear was increased by inhibiting autophagy. To our surprise, these neuronal MNs are released into the extracellular space close to the surface of the cerebral cortex and incorporated into microglia residing in the superficial layer. The micronucleus incorporation influences morphological changes mediated by cGAS. Strikingly, the bulk RNA-seq analyses have revealed that microglia incorporating MNs increase expression level of extracellular matrix (ECM)-related genes. In addition, gene expression signature resembles BAMs. Therefore, our findings demonstrate that MN is a form of intercellular messenger that mediates neuron-to-microglia communications. These results shed light on the novel mechanisms by which microglial functions and states are modulated by neuronal micronuclei in the early postnatal neocortex.

## Results

### Neuronal micronuclei are generated from migrating neurons

Neurons in the cerebral cortex are generated from NPCs in the ventricular and subventricular zone during the embryonic period and migrate to the brain surface. Previously, we observed that MNs exist in the brain of 2-month-old mice^39^. To examine whether MNs also appear in the developing brain, we analyzed the prenatal brain on embryonic day (E) 18.5. We found a couple of MNs in the cerebral cortex (Fig. 1a). Notably, most MNs exist in the superficial layers [marginal zone (MZ) and primitive cortical zone (PCZ)^40^] with few in the cortical plate (CP) (Fig. 1b and 1c). Some of them reside close to nuclei with damaged membranes (Fig. 1d). We next investigated the mechanisms of how MNs are generated in the embryonic brain. Since cancer cells produce MNs via passing a narrow space and deforming nuclei^27, 28^, we examined whether neuronal MNs are also produced by analogous machinery. To test this idea, we introduced an EGFP-expressing plasmid into cells along a lateral ventricle using *in utero* electroporation (IUE) to label migrating neurons^41^. In the E18.5 cortex, EGFP-labeled neurons were squeezed in a PCZ, generating a micronucleus at the tip of the process (Fig. 1e). Next, to analyze the effect of passing a packed neuronal cell-body-rich area, we produced neuronal aggregates by ectopic expression of Reelin^42^. Consequently, several neurons passing aggregates contained MNs (Fig.1f). This effect was not dependent on Reelin stimulation *in vivo* and *in vitro* (Extended Data Fig. 1a to 1e), implying that micronuclear formation was associated with migrating stress. Then, we investigated whether deforming nuclei by passing them through a narrow space influences micronuclear formation *in vitro*. To do this, we designed the trans-well migration assay (Extended Data Fig. 1f). Neurons migrated from the top toward the bottom side exhibited characteristic morphology with deformed and herniated nuclei (Extended Data Fig. 1g). In addition, more MNs were detected in neurons that passed through the membrane depending on the pore size. (Extended Data Fig. 1h to 1j). To further examine the effect of physical stress, we stressed neurons by pumping cells with a syringe (Extended Data Fig. 1k). We found that the number of MNs was increased when the cell density and stroke frequency were increased (Extended Data Fig 1l and 1m), supporting our idea that MNs formation are caused by physical stress, like passing a narrow and dense region. Next, we examined whether MNs observed in the embryonic period are also present in the neonatal brain as neuron-derived micronuclei. To do this, we observed *NexCre:LSL-SUN1-sfGFP-Myc* mice (*NexCre:LSL-Sun1-GFP*), which can specifically label the nuclear envelope of excitatory neurons^43, 44^. The GFP-positive (GFP^+^) small nuclei were detected at P0 and P7 in Layer 1. Also, these small nuclei were retained until P14 (Fig. 1g and 1h). Furthermore, neuronal MNs tended to exist in the motor and somatosensory cortex (Fig. 1i and 1j). The data suggest that neuronal micronuclei produced during brain development reside in the cortical surface, especially in the motor and somatosensory cortices.

**Fig. 1.**
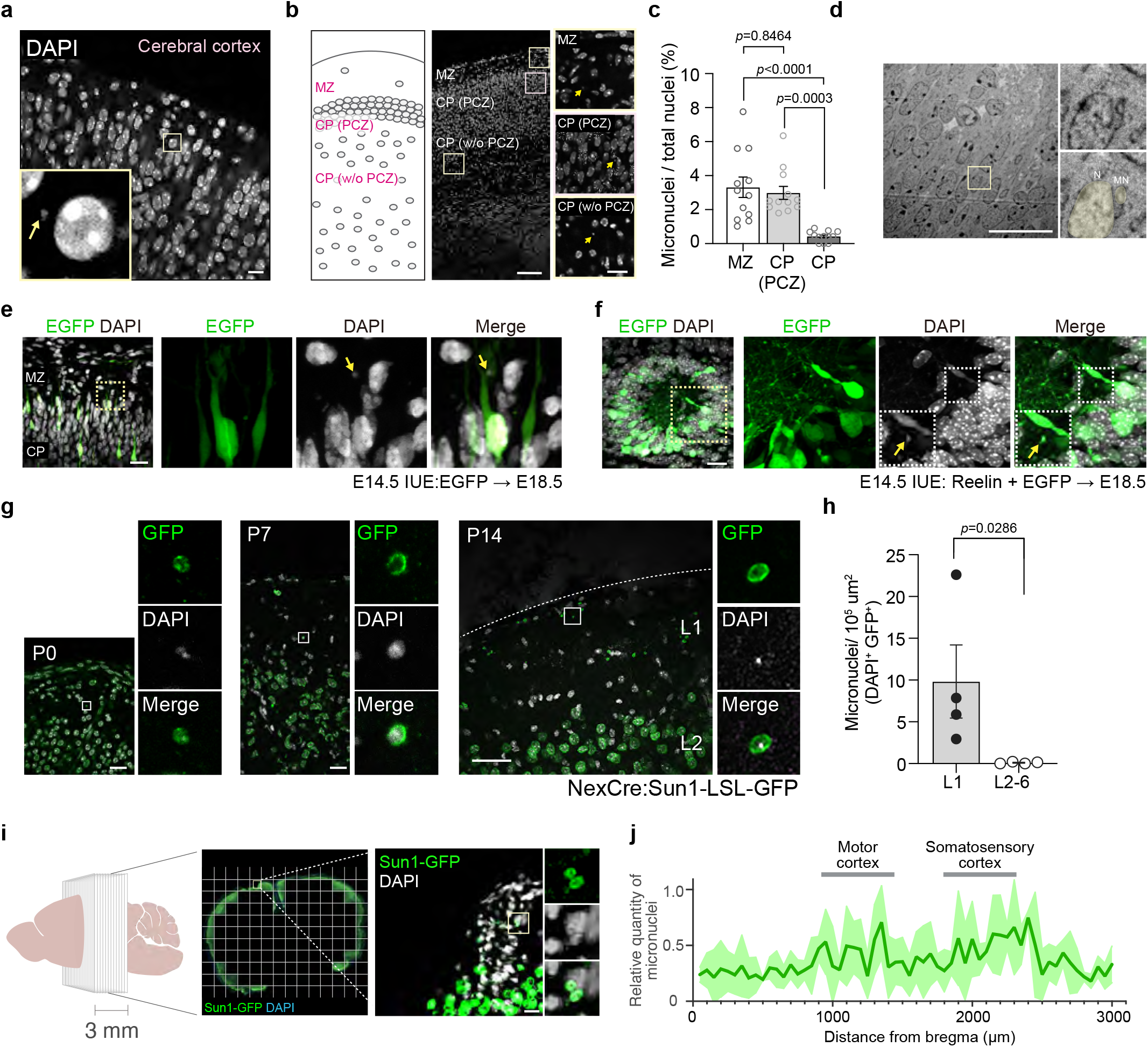
Neuronal micronuclei are generated from migrating neurons. (a) Staining of nuclei in the cerebral cortex of embryonic mice (E18.5). The yellow arrow indicates the micronucleus. Scale bar: 10 μm. (b) MNs mostly exist in the MZ and CP (PCZ) but not CP (w/o PCZ). Scale bar: 100 μm (left), 20 μm (right), E18.5 (c) The graph shows the population of MNs in each field. 12 fields obtained from 2 brains, 2.56 x 10^4^ μm^2^/field; mean ± SEM, *p* values were analyzed by one-way analysis of variance (ANOVA) Tukey’s multiple comparisons test. (d) The transmission electron microscopy (TEM) images of the cerebral cortex of ICR mice. E18.5. N; Nucleus, MN; Micronucleus, Scale bars: 20 μm. (e) Immunostaining of migrating neurons in the cerebral cortex. EGFP plasmids were introduced using *in utero* electroporation at E14.5. The GFP^+^ neurons were detected at E18.5. MN exists in the neuronal process in the vicinity of the brain surface. Yellow arrows indicate MNs. Scale bar: 20 μm. (f) Immunostaining of migrating neurons in the aggregated area. Reelin plasmids were introduced using *in utero* electroporation at E14.5. Scale bar: 20 μm. Yellow arrows indicate micronucleus. The GFP^+^ neurons were detected at E18.5. MNs exist in the processes in the center of neuronal aggregation. (g) Immunostaining of GFP^+^ small nuclei in the cerebral cortex of *NexCre:LSL-Sun1GFP* mice. P0 (left), P7 (center), and P14 (right). Scale bar: 20 μm (P0 and P7) and 50 μm (P14). A dotted line indicates the pial surface. (h) The graph shows the percentage of GFP^+^-MNs in the cerebral cortex. n=4 mice, mean ± SEM, *p* value was analyzed by Mann-Whitney test. (i) Immunostaining of *NexCre:LSL-Sun1GFP* mice brain. The high-magnification images obtained from the cerebral cortex of *NexCre:LSL-Sun1GFP* mice were integrated into a tiling image (center). The typical MNs in Layer 1 are shown in the right panel. P14. Scale bar: 20 μm. (j) The data shows the frequency of micronuclear appearance in Layer 1 of the cerebral cortex. The vertical axis represents the relative value when the maximum number of MNs is defined as 100. The data was quantified using MATLAB software, CAMDi^39^.

### Neuronal micronuclei are propagated to microglia

Since MNs are eliminated by autophagy^45^, we next analyzed micronuclear behavior in wild-type (WT) and excitatory neuron-specific Atg7 conditional knockout mice (*NexCre:Atg7^f/f^*). Atg7 is an E1-like activating enzyme that plays an important role in autophagosome formation. A defect in the *Atg7* gene leads to a delay in autophagosome formation and perturbs the process of autophagy ^46, 47^. The number of neuronal MNs in *NexCre:Atg7^f/f^* brain was increased compared to that in WT (Extended Data Fig. 2a and 2b), suggesting that loss of autophagy activity delays MNs turnover during the postnatal period. Surprisingly, we discovered that several MNs were increased in not only neurons but microglia in *NexCre:Atg7^f/f^* mice even though microglial autophagy was normal (Extended Data Fig. 2c and 2d). Thus, we hypothesized that the neuronal MNs are transferred from neurons to microglia (Fig. 2a). To examine this, we first analyzed whether MNs exist in the extracellular space *in vitro* and *in vivo*. Consequently, several MNs were detected outside primary cultured neurons. In addition, we observed micronuclei in the MAP2^-^ and S100β^-^ regions *in vivo* (Fig. 2b and 2c). Notably, some of the MNs were surrounded by membrane-like structures (Fig. 2d), implying that these MNs are contained in extracellular vesicles.

**Fig. 2.**
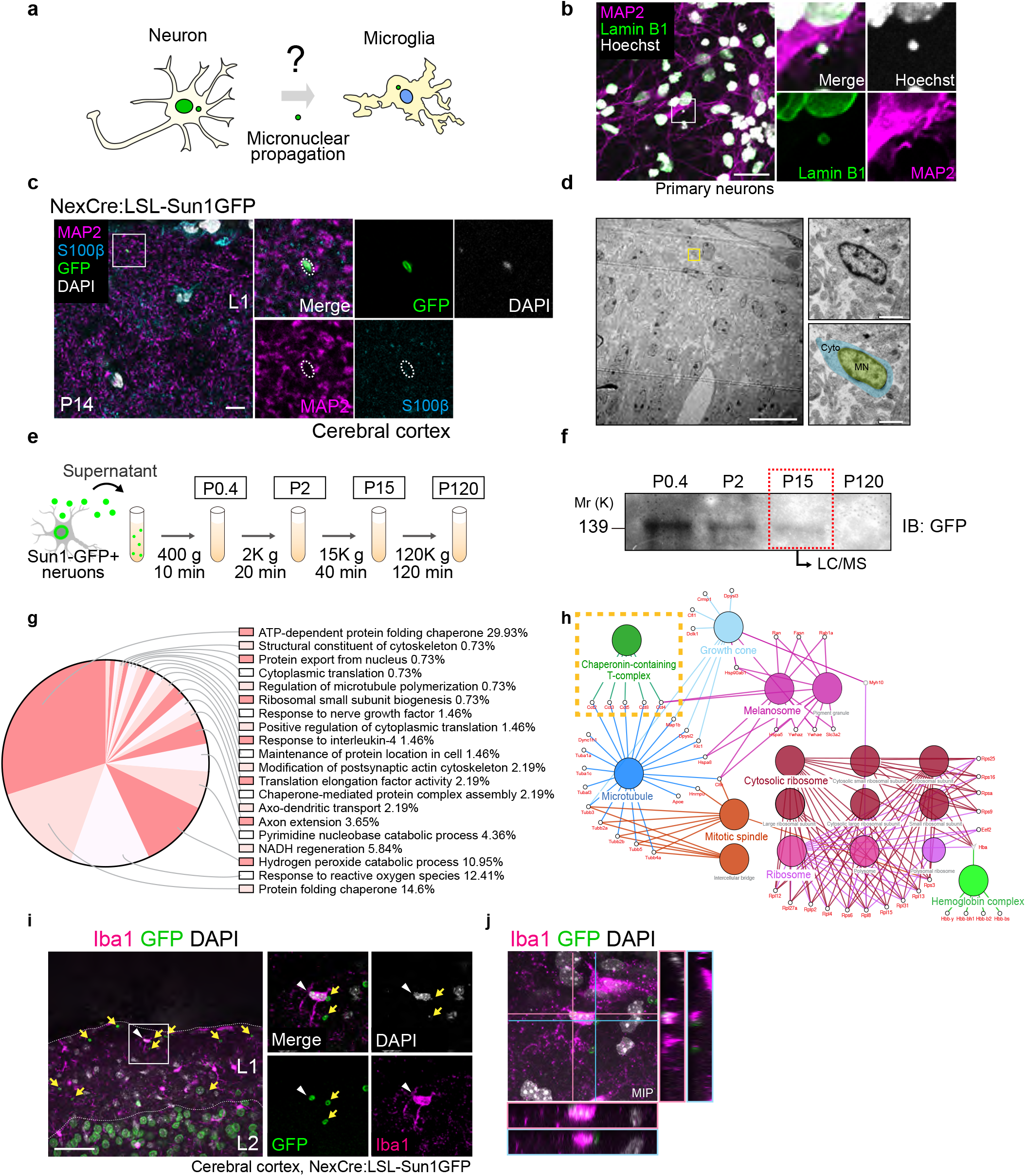
Neuronal micronuclei are propagated to microglia. (a) Schematic illustration of micronuclear transfer from neuron to microglia. (b) Staining of micronucleus in primary cultured neurons (5 DIV). Scale bar: 20 μm. (c) Immunostaining of GFP^+^ small nuclei in the extracellular space (MAP2^-^, S100β^-^) of cerebral cortex. P14. Scale bar: 10 μm. (d) TEM images of the cerebral cortex of ICR mice. E18.5. MN; Micronucleus, Cyto; Cytoplasmic space, Scale bars: 20 μm (left) and 1.0 μm (right). (e) Schematic illustration for fractionation of extracellular MNs. P0.4: apoptotic cells fraction, P2: nuclear fraction, P15: micronuclear fraction, P120: exosome fraction. (f) Immunoblotting of the extracellular MNs obtained from primary cultured cortical neurons of *NexCre:LSL-Sun1GFP* mice. (g) Pie chart showing the composition of the extracellular MNs by gene ontology (GO) analysis. (h) The GO mapping and functional annotation of extracellular MNs components. (i) Immunostaining of GFP^+^-MNs in the Iba1^+^ microglia in Layer 1. *NexCre:LSL-Sun1GFP* mice, P14. Scale bar: 50 μm. White arrowheads indicate MNs in microglia. Yellow arrows indicate extracellular MNs. (j) Image is the inset of Fig. 2i. Neuronal micronucleus is taken in the microglia. MIP: maximum intensity projection image, interval: 1 μm per image, 11 Z-stack images.

Then, to investigate whether MNs are released from neurons, we developed an *in vitro* assay system. First, we biochemically collected MNs from a conditioned medium and identified the fraction containing neuronal MNs (Fig. 2e and 2f). Notably, extracellular MNs were increased when lysosomal activity was inhibited without apoptosis (Extended Data Fig. 2e to 2g), suggesting that a defect in lysosomal activity promotes micronuclear secretion. We next attempted to identify the content of MNs. Using the liquid chromatography and mass spectrometry (LC/MS) analysis, we found several nuclear proteins, such as various Histone proteins (H1.4, H2A type 2-C, H2B type 1-B, H2A.V, H3.3, H1) and Histone-binding protein RBBP7. We also identified several components, including TRiC complex proteins (TCP1α, TCP1β, TCP1δ, TCP1ε, TCP1γ, TCP1ζ, TCP1θ), Tubulin proteins (Tubulin α1A, α1C, αl3, β2A, β2B, β3, β4A, β5), and Rab family proteins (Rab1A, Rab1B, Rab2A, Rab3A, Rab11B, Rab15, Rab35) (Fig. 2g and 2h, Extended Data Table 1). To confirm whether these proteins reside in MNs, we harvested extracellular MNs, adhered to the coverslips, and conducted immunostaining assay. Tubulin and Rab35 signals were observed in the extracellular MN-like structure (Extended Data Fig. 2h and 2i). We also confirmed that TRiC protein complexes reside in MNs in primary cultured neurons using the trans-well analysis (Extended Data Fig. 2j). Moreover, small amounts of Tubulin were colocalized with the neuronal MNs *in vivo* (Extended Data Fig. 2k), suggesting that the proteins identified by LC/MS analysis are potential candidates for the component of MNs. Previously, it has been shown that TRiC complex proteins, TCPβ, serve as a receptor of LC3 and promote autophagosome formation^48^. Because LC3 was localized on the MNs in the cerebral cortex (Extended Data Fig. 2l), the TRiC protein complex may serve as a potential mediator linking MNs to autophagosome formation associated with autophagic secretion (Extended Data Fig. 2m).

Next, we investigated whether extracellular MNs were incorporated into microglia. To examine this, we developed an *in vitro* micronuclear transfer assay system (Extended Data Fig. 3a) and observed the MN-like structure in microglial cell line, BV2 cells (Extended Data Fig. 3b and 3c). Using this system, we analyzed the timing of micronuclear incorporation. BV2 cells took neuronal MNs (GFP^+^-MNs) efficiently at two hours after stimulation with conditioned medium (CM) from *NexCre:LSL-SUN1-sfGFP-Myc* neurons. Whereas the population of MNs in BV2 cells was high at two hours and six hours after stimulation (Extended Data Fig. 3d and 3e), suggesting that microglia take neuronal MNs, followed by disappearing the nuclear envelope or inducing novel MNs derived from microglial nuclei. To verify whether the increase in microglial MNs depends on extracellular MNs, we filtered the CM to remove extracellular MNs. Treatment of filtered CMs did not induce the formation of MNs (Extended Data Fig. 3f and 3g), implying that extracellular MNs are necessary for increasing the number of MNs in microglia. Next, to investigate whether propagation of MNs is observed *in vivo*, we observed *NexCre:LSL-Sun1-GFP* brain. As we expected, most GFP^+^-MNs existed in Layer 1. In addition, some MNs were taken up by microglia (Fig. 2i - 2j). Moreover, we conducted a cortical slice culture imaging using microglia and nuclear labeled mice (*Cx3cr1-EGFP*;*H2B-mCherry* mice)^49, 50^. Intriguingly, several motile MNs were detected, and some seemed to be taken by microglia in the superficial area of cerebral cortex (Extended Data Fig. 3h, Extended Data Video 1). These data are consistent with the possibility that micronuclear propagation regulates microglial characteristics close to the surface of the cerebral cortex.

**Fig. 3.**
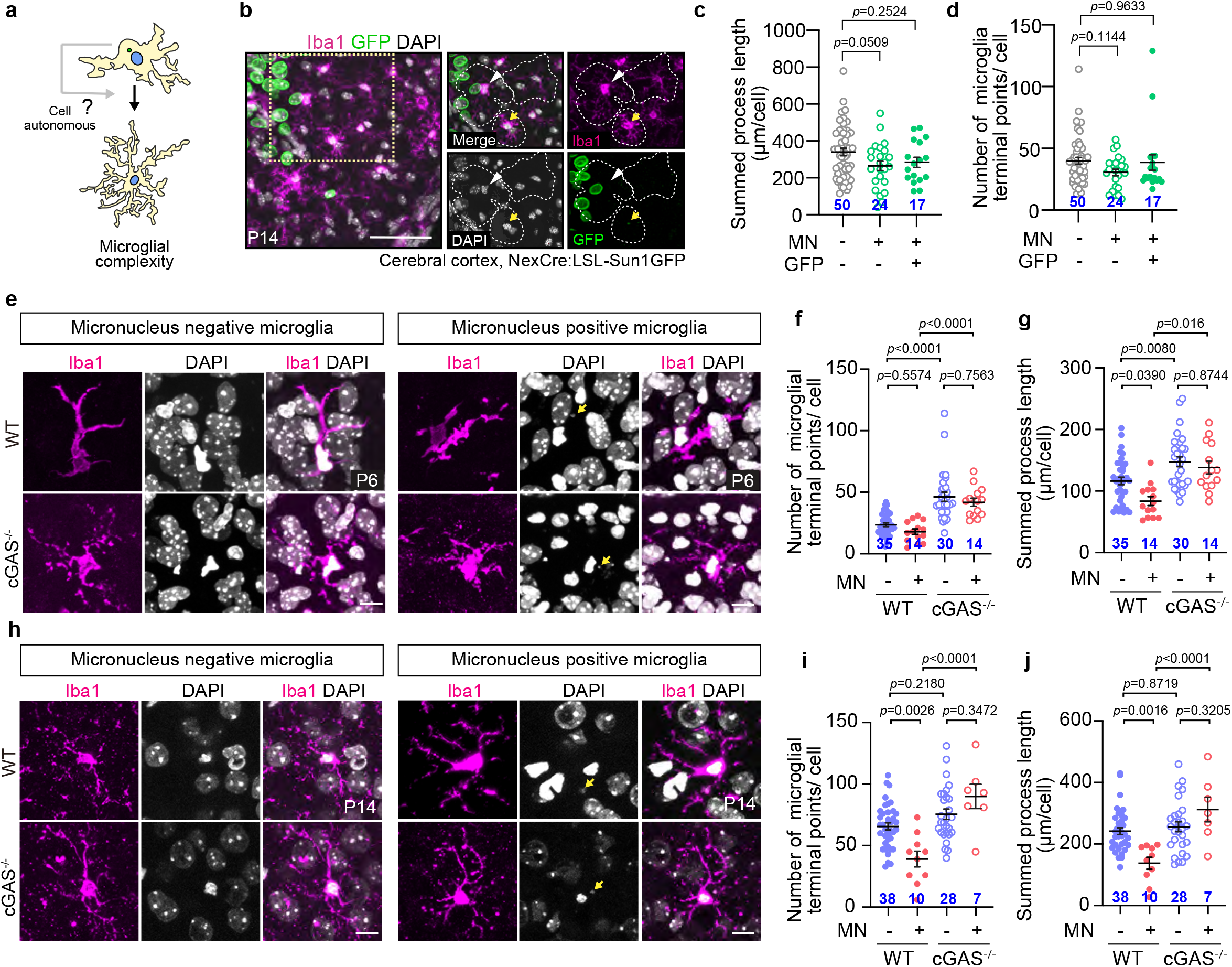
Micronuclei affect microglial morphology during the early-postnatal period via cGAS. (a) Schematic illustration of microglial complexity. (b) Immunostaining of Iba1^+^ microglia with or without neuronal MNs. White arrowheads indicate GFP^-^ micronucleus (GFP^-^-MN). Yellow arrows indicate GFP^+^ micronucleus (GFP^+^-MN). Scale bar: 50 μm. (c) The graph shows the process length in the presence or absence of MNs. 4 brains. mean ± SEM, *p* values were analyzed by one-way ANOVA Dunnett’s multiple comparisons test. (d) The graph shows the number of processes in the presence or absence of MNs. 4 brains. mean ± SEM, *p* values were analyzed by one-way ANOVA Dunnett’s multiple comparisons test. (e) Immunostaining of Iba1^+^ microglia in the cerebral cortex of either WT or *cGAS^-/-^* mice. P6. MN^-^ microglia (left) and MN^+^ microglia (right). Yellow arrows indicate MNs. Scale bar: 10 μm. (f)(g) The graphs show the analyses of microglial morphology. 2 brains. (f) the number of processes, (g) process length, mean ± SEM, *p* values were analyzed by one-way ANOVA Tukey’s multiple comparisons test. (h) Immunostaining of Iba1^+^ microglia in the cerebral cortex of either WT or *cGAS^-/-^* mice. P14. MN^-^ microglia (left) and MN^+^ microglia (right). Yellow arrows indicate MNs. Scale bar: 10 μm. (i)(j) The graphs show the analyses of microglial morphology. 2 brains. (i) the number of processes, (j) process length, mean ± SEM, *p* values were analyzed by one-way ANOVA Tukey’s multiple comparisons test.

### Micronuclei affect microglial morphology during the early-postnatal period via cGAS

To investigate whether MNs regulate microglial characteristics, we first analyzed microglial morphology. The microglia gradually change their shapes and mature by P14^51, 52^. Because MNs were mainly observed in Layer 1, we first examined the involvement in the layer specificity of microglial complexity. Microglia in Layer 1 exhibited slightly less complexity than those in other layers (Extended Data Fig. 4a - 4d), implying a potential link between MNs and microglial complexity. Next, we analyzed whether incorporated MNs affect microglial complexity (Fig. 3a). The process length in microglia is distinguishable according to the existence of MNs, although microglia with GFP^-^-MNs show more simplified morphology (Fig. 3b - 3d). Moreover, there were only minor differences observed in the expression levels of microglial marker genes (*Cd68*, *Tmem119*, *P2ry12*) between cells with and without MN, and no significant biological implications were observed (Extended Data Fig. 5a - 5f). These data suggest that micronuclear propagation changes a hallmark of microglia, such as morphology, regardless of representative microglial gene expression. The question is how MNs regulate microglial states. MNs have been reported to be recognized by cGAS after exposing chromatin DNA^35–38^. Since cGAS is mainly expressed in microglia in the brain^53^ (Extended Data Fig. 6a), we analyzed conventional *cGAS^-/-^* mice brains. The morphological complexity was increased in *cGAS^-/-^* microglia at the early stage (Extended Data Fig. 6b - 6e), implying that cGAS is crucial for regulating morphological complexity. Notably, cGAS deficiency cancels the effect of MNs-regulated morphological changes (Fig. 3e - 3j, Extended Data Fig. 6f - 6i). These data suggest that cGAS is involved in MN-dependent morphological changes.

**Fig. 4.**
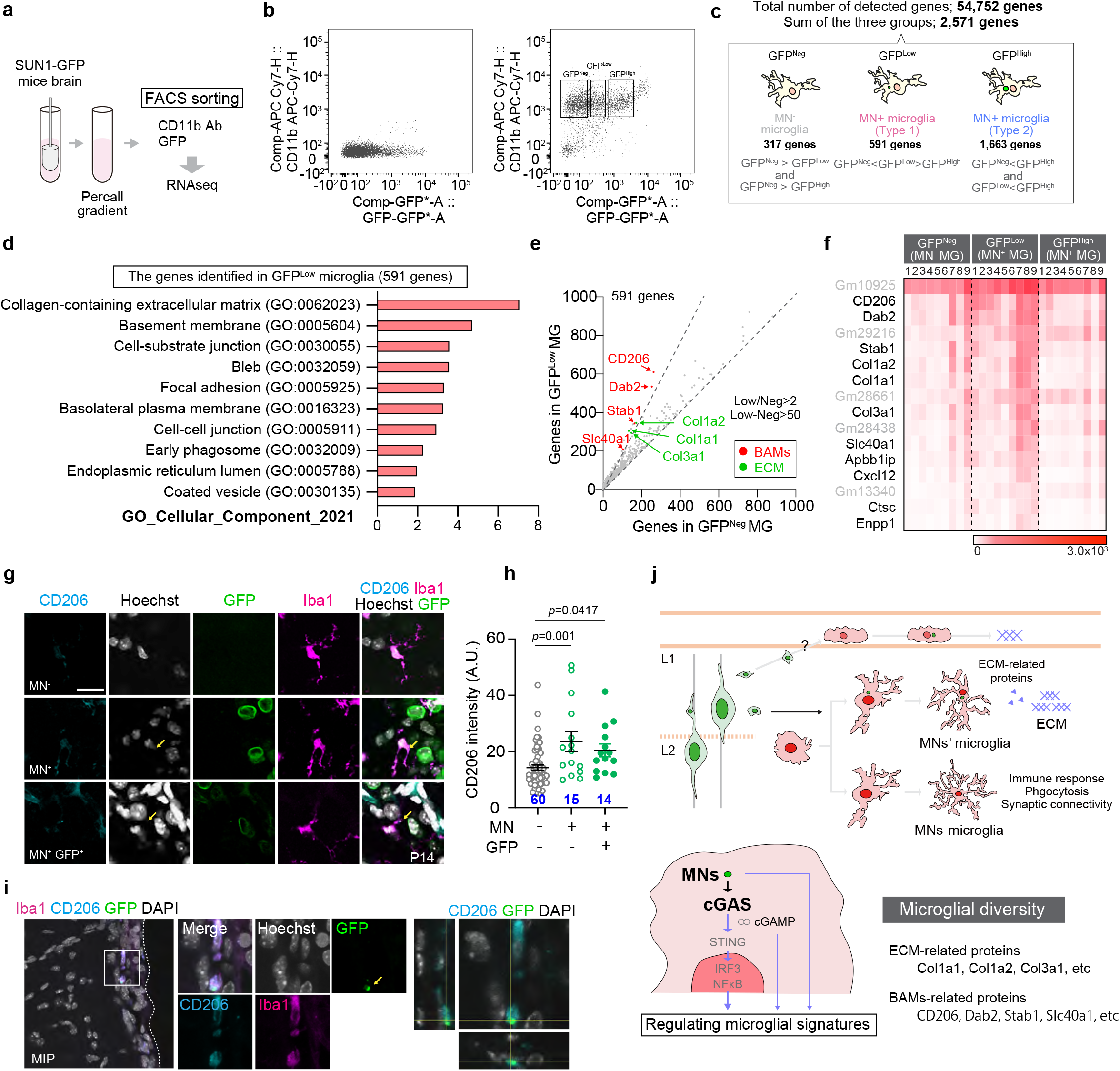
Neural micronuclei alter a hallmark of microglia. (a) The *NexCre:LSL-Sun1GFP* mice brains were homogenized using Dounce tissue homogenizer. Myelinated materials were removed using Percoll gradient. Microglia were purified by FACS using anti-CD11b antibody and GFP signals, followed by conducting bulk-RNA seq analysis. (b) Representative gating strategy for flow cytometry of either GFP^+^ or GFP^-^ microglia from the *NexCre:LSL-Sun1GFP* whole brain (P8, P9, P10). (left) Non-microglial population, (right) Microglial population, left gate; GFP^Neg^ microglia, center gate; GFP^Low^ microglia, right gate; GFP^High^ microglia. (c) The simplified diagram for the deferential gene expression (DEG) analysis. The number of DEG in each microglial population categorized by GFP intensity (GFP^Neg^, GFP^Low^, GFP^High^). (d) The GO enrichment analysis of DEG in GFP^Low^ MN^+^ microglia (591 genes). Data was analyzed in Enrichr analysis tool. (e) The scatter plot indicates the ratio and differences of FPKM between GFP^Neg^ MN^-^ microglia and GFP^Low^ MN^+^ microglia. FPKM: GFP^Low^/GFP^Neg^>2, GFP^Low^-GFP^Neg^>50. (f) The Heat map indicates the comparison of FPKM among GFP^Neg^ MN^-^ microglia, GFP^Low^ MN^+^ microglia, and GFP^High^ MN^+^ microglia. FPKM: GFP^Low^/GFP^Neg^>2, GFP^Low^-GFP^Neg^>50. (g) Immunostaining of Iba1^+^ microglia with or without neuronal MNs. White arrowheads indicate GFP^-^ micronucleus (GFP^-^-MN). Yellow arrows indicate GFP^+^ micronucleus (GFP^+^-MN). Scale bar: 50 μm. (h) The graph shows the fluorescence intensity of CD206 in the presence or absence of MNs. More than 13 images obtained from two independent *NexCre:LSL-Sun1GFP* brain, P7; mean ± SEM, *p* values were analyzed by one-way ANOVA Dunnett’s multiple comparisons test. (i) Immunostaining of Iba1^+^ microglia with neuronal MNs in the meninges. Yellow arrows indicate GFP^+^MN. (j) Schematic model for regulating microglial state by neuronal MNs.

### Neuronal micronuclei alter a hallmark of microglia

Because neuronal MNs seem to affect the microglial state, we purified CD11b^+^ GFP^±^ microglia from the postnatal brains of *NexCre:LSL-Sun1-GFP* mice. The mRNAs isolated from microglia were analyzed by bulk RNA-sequencing (RNA-seq) analysis, and the differentially expressed genes (DEG) were determined using *edgeR* software^54, 55^ (Fig. 4a and 4b, Extended Data Fig. 7a, Extended Data Table 2). Due to the presence of SUN1-GFP in MN, we first divided the microglial subpopulation into two groups [GFP^-^ microglia and GFP^+^ microglia]. Since the population of GFP^+^ microglia was not converged, we further subdivided GFP^+^ microglia into two groups; Type 1 (low signal of GFP, GFP^Low^) and Type 2 (high signal of GFP, GFP^High^) (Fig. 4c). RNA-seq analysis revealed that GFP^High^ microglia possessed various neuronal mRNA (*MAP1b*, *MAP2*, *Rian*, *Grin2b*, *Pgm2l1*) (Extended Data Fig. 7b - 7r). Since phagocytic microglia often possess mRNAs of engulfed cells^56^, we speculated that neuronal mRNA in GFP^High^ microglia are derived from phagocytosed apoptotic neurons. In contrast, as little neuronal mRNA was detected in GFP^Low^ microglia, we analyzed these GFP^Low^ microglia as microglia incorporated neuronal MNs. Intriguingly, GFP^Low^ microglia increased the expression of ECM-related genes compared to both GFP^Neg^ and GFP^High^ microglia (Fig. 4d). The genes identified in GFP^Low^ microglia were partially shared with the genes which were highly expressed in the BAMs (*CD206*, *Dab2*, *Stab1*, *Slc40a1*) ^16, 57, 58^ (Fig. 4e and 4f, and Extended Data Fig. 8a - 8v), suggesting that neuronal MNs are involved in changing microglial state. Indeed, some microglia with MNs slightly increased the expression of CD206 (Fig. 4g and 4h). Moreover, we found BAMs in the meninges also incorporated neuronal MNs with few chromatin (Fig. 4i). These data suggest that neuronal MNs regulated-microglial signature may be controlled by multiple pathways.

## Discussion

So far, MNs have been described mainly in the context of tumorigenesis, in which they emerge by genomic instability and cellular migration. DNA fragments of the MNs are released to the cytoplasm, and cell-autonomously accelerate the malignancy phenotype^59^. In this study, however, we have revealed the existence of MNs in a physiological context and their role in the developmental regulation of microglia. Remarkably, our results demonstrated that MNs generated in neurons are released into the extracellular space and incorporated into microglia, regulating microglial signatures (Fig. 4j). Our study thus unexpectedly unveils a novel form of intercellular communication via MNs.

The finding in this study is that incorporating MNs into microglia changes the properties of those microglia. Notably, the ECM-related genes were upregulated in MNs^+^ microglia. Furthermore, the gene expression profile of MNs^+^ microglia partially overlapped with that of BAMs. Therefore, we hypothesized that neuronal MNs might endow microglia with properties similar to BAMs. BAMs reside in the meninges, perivascular spaces, and choroid plexus. Fate mapping analyses have revealed that EMPs from the yolk sac differentiate into maturating macrophages^60^. Then, they change into either parenchymal microglia or BAMs depending on whether EMPs are stimulated by TGFβ signaling^16^. However, understanding the functions of BAMs that underlie brain development and homeostasis has been hindered due to a low number of cells. On the other hand, despite the limited number of studies, several groups have reported the biological relevance of BAMs. First, ablating BAMs using pharmacological inhibitors increased aberrant ECM accumulation^61^. Indeed, the BAMs-residing areas, such as meninges, choroid plexus, and perivascular spaces, are surrounded by various ECM proteins. Thus, BAMs may regulate the architecture of the border region through the precise expression of ECM-related proteins. Perivascular macrophages have been reported to repair a small lesion in the vessel, suggesting that BAMs contribute to maintaining and repairing the brain vessels^62^. Here, we found that microglia taken up neuronal MNs change ECM-related gene expressions (*Col1a1*, *Col1a2*, *Col3a1*), implying that extracellular MNs are one of the potential mediators that alters the microglial signature during early-postnatal stages. Because the meningeal macrophages seem to transit to the perivascular macrophages during the postnatal stages^63^, it is likely that microglia or meningeal macrophages taken up MNs migrate toward the perivascular region. We also observed that various neuron-specific genes are expressed in GFP^High^ microglia. Since early postnatal microglia are mostly reactive states and maintain the brain environment through phagocytosing apoptotic neurons, it is likely that GFP^High^ microglia may also compose GFP^+^ neuronal nuclei derived from apoptotic neurons. Meanwhile, the genes identified in RNA-seq analysis, such as Nrgn and Marcksl1, are expressed in non-neuronal cells, including microglia. So, we presume that some may be target genes induced by MNs.

The mechanisms that underlie the release of MNs have not been clarified. While the autophagy pathway mainly plays a vital role in cargo degradation, either acceleration or inhibition of autophagic machinery modulates the secretion pathway^64^. In this study, we found a couple of potential targets that underlie MNs secretion associated with autophagy, such as TCP family proteins and Rab family proteins. It has been shown that treatment with bafilomycin, which can inhibit autophagosome-lysosome fusion via perturbed pH in the lysosome, promotes aggregated α-synuclein secretion^65^. Because TRiC protein subunit, TCPβ, bridges between aggregated protein and LC3^48^, it is possible that MNs are also secreted by analogous machinery mediated by TCP subunits. Also, Rab family proteins, Rab11 and Rab35, have been reported to be involved in regulating exosome secretion^66, 67^. Since we identified these proteins as potential components in our proteomics screening, these Rab family proteins may also serve as regulatory factors that control MNs secretion. The ATG-conjugation system plays a critical role in the degradation of the inner autophagosomal membrane but not in the formation of an autolysosomes^68^. While we cannot completely rule out the possibility that the extracellular vesicles are apoptotic bodies formed by dying neurons, we speculate that the extracellular vesicles were switched from the autophagy degradation pathway to the secretory pathway and subsequently released due to a loss of autophagy function (see Extended Data Fig. 2m).

To date, EVs, including microvesicles, have been identified as key mediators of intercellular communication^69^. Microvesicles are similar in size to extracellular MNs (1.0 μm to 3.0 μm), and are released from the plasma membrane via a budding process, transferring cargo to target cells. The ESCRT III complex is crucial for the scission of vesicles from the membrane^70^ and regulates membrane repair by releasing the damaged patch to the extracellular region^71, 72^. Thus, it is likely that the ESCRT-III complex may control extracellular MNs production through the budding process, which is linked to the autophagy-dependent pathway. Intriguingly, EVs transport organelle intercellularly in the brain. Mitochondria are among the organelles that are transported by EVs under both neurodegenerative or ischemic conditions^73, 74^. Hence, EVs containing MNs may represent a novel type of microvesicle that modulate microglial signature. In addition to microvesicles, various types of EVs, including exosomes, have been reported to contribute to cell-cell communication in the central nervous system^69^. Exosomes are small vesicles (ranging from 100 nm to 1.0 μm) that are released from the multivesicular bodies and carry both RNAs and proteins to target cells. Under pathogenic conditions, glia propagate the pathogenic proteins through exosomes^75^. Components of the nucleus, such as nuclear protein (Lamin A/C and Histone H2B) and chromatin, are transferred from MNs to exosomes and released to the extracellular space^76^. Although the size of MNs is larger than the exosome, a comparative analysis between the exosome and microvesicle is an exciting study, and further validation is required.

Recent research suggests that cGAS is a redundant DNA sensor in the cytoplasm^77^. Previous studies have shown that cGAS in microglia is involved in ischemia, virus infection, and neurodegeneration^78^. Since cGAS induces IFNβ expression by activating the cGAS-STING-IRF3 pathway, we investigated the link between incorporating MNs and IFNβ expression. However, we did not obtain clear data with or without MNs *in vivo*. While we cannot rule out the possibility of technical difficulties to detect endogenous IFNβ, we also speculate an IRF3-independent mechanism. Previous studies have reported that cGAMP, an intercellular mediator, is transported via specific transporters^79–81^. Hence, it is intriguing that released cGAMP regulates microglial characteristics via an autocrine pathway. Furthermore, as we observed that certain MNs lack chromatins, it is likely that the MN-dependent, cGAS-independent pathway is also involved in regulating microglial signature.

During embryonic development, microglia are believed to migrate to the pial surface in response to signals from meninges^82^. As the microglial signature is region- and stage-specific, it is possible that MNs in the superficial layer contribute to generating the unique microglial signature. In the cerebral neocortex, neurons form six-layer structures. In the PCZ, a dense region in the superficial CP, migrating neurons contain intracellular MNs with deformed nuclei^40^. We often observed here that intracellular MNs reside in the process of migrating neurons with deformed nuclei in the PCZ. Thus, we hypothesize that physical stress may trigger the formation of intracellular MNs in neurons, as the number of MNs^+^ neurons increases when neurons pass through narrow spaces *in vitro*. Similar to cancer cells^27, 28^, migrating neurons may receive physical stress and nuclear deformation, which can induce changes in their epigenetic state^83^. Therefore, it is likely that physical stress in the neuronal nucleus provides a biological cue that affects the gene expression profile associated with the microglial states.

In conclusion, our study demonstrates that MNs are generated from migrating neurons and released during the early postnatal period. These MNs are subsequently taken up by microglia, and their presence is associated with cGAS activity and alterations in the microglial state. Our findings provide novel insights into the relevance of the propagation of neuronal MNs and microglial diversity during early-postnatal stages.

## Materials and Methods

### Mice

The Crl:CD1(ICR) mice were obtained from Japan SLC, Inc. (Shizuoka Japan). The B6.Cg-Atg7^<tm1Tchi>^ (Atg7^f/f^) mice have been described previously^46^. The B6;129-Gt(ROSA)26Sor^tm5(CAG–Sun1/sfGFP)Nat/J^ (*LSL-Sun1-GFP*), B6.Cg-Gt(ROSA)26Sor^tm9(CAG-tdTomato)Hze/J^), B6(C)-cGAS^tm1d(EUCOMM)Hmgu/J^ (*cGAS*^*-/-*^), and B6.129P2(Cg)-Cx3cr1^tm1Litt^/J mice were obtained from the Jackson Laboratory (the stock #: 021039, 007909, 026554, and 005582 respectively)^43, 44, 49, 84^. R26-H2B-mCherry (accession #: CDB0239K) mice were previously described^50, 85^. To generate *NexCre:Atg7^f/f^* mice and *NexCre:LSL-Sun1-GFP* mice, the *Atg7^f/f^* mice and *LSL-Sun1-GFP* mice were crossed to the Neurod6^tm1(cre)Kan^ (NexCre)^86^, respectively. All of the animal experiments were conducted according to the university guidelines for animal care. This study was approved by the Animal Experiment Committee in the University of Tsukuba (the approval numbers:18-257, 19-340, 20-438, 21-443, 22-427).

### Cell culture

BV2 cells were cultured in DMEM/nutrient mixture F-12 (DMEM/F12; Thermo Fisher Scientific) containing 10% fetal bovine serum (FBS), 100 units penicillin, and 100 mg streptomycin (P/S; Thermo Fisher Scientific) at 37LJ with 5% CO ^87^. Neuro2A cells were cultured in EMEM (FUJIFILM Wako) containing 10% FBS and P/S. HEK293T cells were cultured in DMEM (FUJIFILM Wako) containing 5% FBS and P/S^88^. Primary cortical neurons were prepared from E13.5-14.5 ICR mice or *NexCre:LSL-Sun1-GFP* mice as previously described^89^. Briefly, mouse cortical neurons were plated on coverslips coated with either 0.01875% Poly (ethylenimine) solution (SIGMA) and cultured in Neurobasal medium (Thermo Fisher Scientific) containing B-27 supplement (Thermo Fisher Scientific) and P/S. Neurons were maintained for 5 - 7 days at 37LJ with 5% CO_2_ and were subjected to *in vitro* MNs transfer assay and trans-well migration assay.

### Antibodies, materials, and plasmids

For immunofluorescence, anti-MAP2 (MAB378, Merck Millipore, 1:500), anti-Iba1 (#019-19741, WAKO, 1:1000), anti-GFP (#598, MBL, 1:500), anti-GFP (ab13970, abcam, 1:1000), anti-RFP (#600-401-379, Rockland, 1:3000), anti-Cleaved Caspase-3 (Asp175) (5A1E, CST, 1:500), anti-CD68 (FA-11, Biolegend, 1:500), anti-Lamin B1 (ab16048, abcam, 1:500), anti-S100β (ab52642, abcam, 1:1000), anti-Rab35 (11329-2-AP, Proteintech, 1:1000), anti-Tubulin (DM1A, SIGMA, 1:1000), anti-LC3 (PM036, MBL, 1:1000), anti-CD206 (AF2535, R&D systems, 1:200), anti-P2ry12 (AS-55043A, AnaSpec, 1:1000), anti-Tmem119 (ab209064, abcam, 1:600) were used as the primary antibody. For immunoblotting, anti-GFP (#598, MBL, 1:1000), anti-Tubulin (SIGMA, DM1A, 1:1000), and anti-Lamin B1 (ab16048, abcam, 1:1000) were used as the primary antibody. Bafilomycin A1 (BafA) was purchased from CST (#54645). The recombinant Reelin (3820-MR/CF) was purchased from R&D Systems. The full-length mouse Reelin cDNA and pCAGGS1 plasmid containing a CAG promoter were kindly provided by Dr. T. Curran (Children’s Mercy Kansas City) and Dr. J Miyazaki (Osaka University), respectively.

### *In utero* electroporation

An *in utero* electroporation was modified as previously described^90^. Briefly, the day detected virginal plug was defined as E0, and the day of birth was defined as P0. The pregnant ICR or C57BL/6 mice (E14.5) were anesthetized with isoflurane. The mother mouse was warmed using a heater to maintain body temperature. After a midline laparotomy was performed, the embryos were pulled out. The 1.0 μl of DNA (5 µg/µl in water) together with the 0.01% Dye Fast Green (SIGMA) was injected through the uterine wall into one of the lateral ventricles of the cerebral cortex by the glass capillary. For injection, the glass capillaries were made from the thin-walled glass capillary (GD-1, Narishige) utilized by the capillary puller (PC-100, Narishige). An *in utero* electroporation was performed using an electroporator (NEPA21, NEPAGENE). The electronic pulse (poring pulse 50V, pulse length 50 msec, pulse interval 950 msec, 4 times and transfer pulse 50V, pulse length 50 msec, pulse interval 950 msec, 4 times) were applied with a forceps-type electrode (CUY650P5, NEPAGENE). The plasmids of pGAG-EGFP and pCAG-Reelin were injected in one side to minimize the damage to the brains.

### Immunohistochemistry

The brains were fixed by perfused with 4% paraformaldehyde (PFA) in phosphate buffered saline (PBS) and fixed with the same fixative solution overnight. After 30% sucrose in PBS infiltration, the samples were embedded in Tissue-Tek OTC compound (SAKURA) and sliced at a 50 μm thickness by the cryostat (Leica Biosystems). Free-floating slices were permeabilized and blocked with PBS containing 0.25% Triton X-100 and 5% bovine serum albumin (BSA). Primary antibodies were incubated in PBS containing 0.1% Triton X-100 and 5% BSA for 3 days (anti-MAP2 antibody) or 1 day (all others) at 4°C. Immunostaining was detected using Alexa 488, Alexa 594, or Alexa 647 fluorescent secondary antibody (Thermo Fisher Scientific, 1:1000), incubated in PBS containing 5% BSA for 1 hour at room temperature. Slices were counterstained with DAPI solution (Dojindo) and mounted in VECTASHIELD Mounting Medium (Vector Laboratories). Tissue specimens were observed using the confocal laser scanning microscope (LSM700, Carl Zeiss) with 20x (Plan-Apochromat 20x/0.8 M27) and 40x (Plan-Apochromat 40x/1.3 Oil DIC M27) objective. The diode excitation lasers (Diode 405, Diode 488, and Diode 555) were operated and directed to a photomultiplier tube (LSM T-PMT, Carl Zeiss) through a series of bandpass filters (Ch1:BP420-475 + BP500-610, Ch2:BP490-635, and Ch3:BP585-1000). The Z-stack images (interval: 1 μm per image) were acquired using ZEN software (Carl Zeiss). The microglial morphological were assessed by the Image J skeleton and fractal analysis^91^. For the high throughput imaging in Figure 1i, the fluorescence images were obtained using the Thunder imaging systems (Leica) equipped with 20x objective (HC PL Fluotar L 20x/0.4 CORR PH1), the diode excitation laser (395 and 470), and filter set (Ex391/32, DC 415, Em 435/30 for DAPI, Ex479/33, DC 500, Em519/25). The fluorescent images were decoded by Thunder imaging system for computational clearing (Leica Application Suite X 3.7.0)

### Immunocytochemistry

Primary cortical neurons were plated at 1.0 x 10^6^ cells/well on the trans-well (Corning, 12 mm Trans-well with 3.0 µm Pore Polycarbonate Membrane Insert, 3402) for 3 days. The polycarbonate membranes (top and bottom) were coated with 0.01875% Poly (ethylenimine) solution (SIGMA) for 1 day. Only the bottom sides were recoated by 1.42 μg/ml laminin (SIGMA) for 1 day. Neurons were stained with anti-MAP2 and anti-Lamin B1 antibodies and subjected to analyzing MNs using MATLAB-based program for quantifying MNs (Calculating automatic MNs distinction: CAMDi)^39^. Neuro2A cells (5.0 x 10^4^ cells) were loaded into a syringe fitted with a needle [Dentronics, No.32 (0.26 mm x 12 mm), Cat# HS-2739B] and were subject to single or ten strokes. Neuro2A cells were plated on the cover glass and were incubated for one day. Cells were then stained with Hoechst and subjected to analyzing MNs using CAMDi. BV2 cells were plated at 2.5 × 10^4^ cell/cm^2^ density on glass coverslips (15 mm diameter, Matsunami Glass) in 12 well plates. Conditioned medium was added 1 day after plating. Cells were fixed with 4% PFA in PBS for 10 min at room temperature. BV2 cells on coverslips were blocked in 0.25% Triton X-100 in blocking solution (5% BSA in PBS) for 30 min at room temperature and then incubated with primary antibodies diluted in blocking solution for overnight at 4°C. After washing with PBS, the cells were incubated with Alexa Fluor Plus secondary antibodies (Thermo Fisher Scientific, 1:1000) or Dylight Fluorochrome conjugated secondary antibodies (abcam, 1:1000) diluted in blocking solution for 30 minutes at room temperature. The nuclei were stained with 10 μg/ml Hoechst 33342 (SIGMA) or 1.0 μg/ml DAPI. The coverslips were mounted onto slide glasses using FLUOROSHIELD Mounting Medium (ImmunoBioScience). Fluorescent images were obtained using the fluorescence microscope (BZ-9000, Keyence) with 40x (PlanApo 40x/0.95) and 100x (PlanApo VC 100x/1.4 Oil) objective and a charge-coupled device camera (Keyence). A proper wavelength was selected using fluorescence filter cubes;DAPI [Ex 360/40, DM 400, BA 460/50] (Keyence), GFP-B [Ex 470/40, DM 495, BA 535/50] (Keyence), mCherry-A [Ex FF01-562/40-25, Di

FF593-Di02-25x36, Em FF01641/75-25](Opto-Line, Inc). Other fluorescent images were obtained using the confocal laser scanning microscope (LSM700, Carl Zeiss) with 20x (Plan-Apochromat 20x/0.8 M27) and 40x (Plan-Apochromat 40x/1.3 Oil DIC M27) objective. The diode excitation lasers (Diode 405, Diode 488, and Diode 555) were operated and directed to a photomultiplier tube (LSM T-PMT, Carl Zeiss) through a series of bandpass filters (Ch1:BP420-475 + BP500-610, Ch2:BP490-635, and Ch3:BP585-1000).

### Immunoblot analysis

To characterize the extracellular MNs released from neurons, the cultured media were collected from primary cortical neurons of *NexCre:LSL-Sun1-GFP* mice or ICR mice, and isolated by differential centrifugation. The collected media was subjected to a centrifugation step of 400 x g for 10 min at room temperature to remove cells. Next, the supernatant was centrifuged at 2,000 x g for 20 min at 4LJ to remove apoptotic bodies. For the collection of large factors, the supernatant was centrifuged at 15,000 x g for 40 min at 4LJ. This supernatant was next subjected to ultracentrifugation at 120,000 x g for 2 h at 4LJ in a SW 41 Ti Rotor Swinging-Bucket rotor (Beckman Coulter), and the pellets collected as small factors like exosome. Cellular proteins and extracellular proteins from media were extracted with 100 μl of ice-cold RIPA lysis buffer (10 mM Tris-HCl [pH7.5], 150 mM NaCl, 1 mM EDTA, 1% NP-40, 0.1% sodium deoxycholate, 0.1% SDS, and 1 µM dithiothreitol [DTT]) and 30 μl of that, respectively. After SDS-polyacrylamide gel electrophoresis, each sample was transferred to a polyvinylidene difluoride membrane (Pall). The membranes were blocked in 5% skim milk for 60 min at room temperature and were then incubated overnight at 4°C with the primary antibodies. After washing with PBS, the membranes were incubated for 1 hour with the peroxidase-conjugated secondary antibodies. The signals on the membrane were detected by Chemi-Lumi One Super (Nacalai tesque)

### *In vitro* micronuclei transfer assay

Primary cultured cortical neurons purified from E13.5-14.5 ICR mice embryonic cerebral cortex were plated at 1.0 x 10^6^ cells/well (12-well plate) and incubated for 7 days and were then incubated with DMSO or 300 nM BafA at 37°C for 3 h. Neurons were rinsed with PBS twice and refreshed with Neurobasal medium containing B27. After a 3 h incubation, the dead cells were removed using centrifugation (pelleted at 800LJ× g, 5 min, 4LJ). To remove MNs from the conditioned medium, the medium was filtrated through 0.22 μm pore PES filters (Millipore).

BV2 cells were treated with the conditioned medium for 2 - 6 hours and subjected to an *in vitro* MNs assay. HEK293T cells were plated at 1.0x10^6^ cells/ dish (6 cm dish) and incubate for 1 day. Cells were then transfected with either Reelin-HA or control plasmid using 1.0 mg/ml polyethyleneimine MAX (Polyscience). After 2 days, the conditioned medium was collected and were centrifuged at 8,000 rpm for 5 min to remove the debris. Primary cultured cortical neurons [5 day in vitro (DIV)]were treated with the conditioned medium for 6 hours and were then subjected to a MNs assay.

### Micronuclei analysis

FIJI ImageJ software (NIH) and ZEN (Carl Zeiss) are used for converting each Z-stack image (interval; 1 μm per image, a total thickness of images staining with MAP2; 10 to 15 μm, a total thickness of images staining with Iba1; 30 to 50 μm) to a single TIFF file. The data were imported into the MATLAB-based program CAMDi for quantifying MNs^39^. Briefly, the rate of overlap area of the nucleus is defined as 0.5. The imported data are conducted binarization and are determined the threshold of each color. After making the binary images, the merged areas are extracted according to the threshold, followed by quantification of the number, area, and volume. It has been reported that the maximum size of micronucleus is defined as one-third of the main nucleus^92, 93^. The major axis of neuronal nuclei is approximately 12 to 15 μm. On the other hand, we rarely observed large sizes of nuclei (approximately less than 18 μm), but they resided in the brain. Thus, the extra nuclei within a major radius of 3.0 μm were defined as MNs.

### Transmission electron microscope

For the cellular electron micrograph, BV2 cells were collected after conducting a micronuclear transfer assay (see [*in vitro* micronuclear transfer assay] section) and subjected TEM imaging. Briefly, cells were fixed with 2.5% glutaraldehyde in 0.1M phosphate buffer at 4°C for 30 min. After washing with phosphate buffer, cells were fixed with 1.0% osmium tetroxide 4°C for 90 min. Next, cells were dehydrated with ethanol and were then reacted with propylene oxide for 20 min twice. After obtaining the thin section, the samples were stained with 2% uranyl acetate and lead citrate. The images were obtained using a JEM 1400 (JOEL). For the brain electron micrograph, the postnatal ICR mice (P0) were perfused with fixative solution (4% PFA, 2.5% glutaraldehyde, 0.1% picric acid, 0.05 mg/ml ruthenium red in 0.1M phosphate buffer [7.2]). The mice brains were post-fixed for 1-2 hours at 4°C using a fixative solution. After washing with phosphate buffer, the brain sections (coronal section, 300 μm) were prepared with a vibratome. The sections were then fixed again with 1% osmium tetroxide and 0.05% ruthenium red for 30 min at room temperature. After sequential dehydration with ethanol solutions and replacement with n-butyl glycidyl ether, the extra-thin sections (60-80 nm) were obtained using ultra cut UCT (Leica) and stained with 2% uranyl acetate and lead citrate. The images were obtained using a JEM 1400 (JEOL).

### Mass spectrometry and data analysis

The extracellular MNs were collected by ultracentrifugation (see the [immunoblot analysis] section). After adding 90 μl of AB buffer (100 mM ammonium bicarbonate buffer [90%]/Acetonitrile [10%]), the samples were added 4 µl of 100 mM DTT/10% acetonitrile/100 mM AB buffer and incubated at 37°C for 60 min. The samples were added 10 μl of 100mM Iodoacetamide/10% Acetonitrile/100mM AB buffer and incubated at 37°C for 30 min. The samples were then subjected to trypsin digestion and injections into LC/MS (Zaplous ADV Q-Exactive system, AMR Inc.). The raw data were categorized using the DAVID Bioinformatics Resources 6.8 database (https://david.ncifcrf.gov/)

### FACS analysis

Isolation of GFP^Neg^, GFP^Low^, and GFP^High^ microglia were performed as previously described^52, 94^. The cortices of the *NexCre:LSL-Sun1-GFP* mice (P8 to P10) was homogenized using a syringe with 23G and 27G needles, sequentially in 500 μL of PBS including 0.2% DNaseI (Takara). The solution was mixed with 145 μL of Debris Removal Solution (Miltenyi), and 600 μL of PBS was layered on it. The tube was centrifuged at 3,000 x g for 10 min at 4°C. After removing upper layer including debris, the solution was mixed with 600 μL of PBS and centrifuged at 1,000 x g for 10 min at 4°C. After removing supernatant, the pellet was resuspended with 200 μL of PBS including 0.2% BSA, added anti-CD11b conjugated with APC-Cy7 (BioLegend, #101226) and anti-CD45 conjugated with PerCP-Cy5.5 (BioLegend, #109828), and placed on ice for 10 min. The solution was mixed with 60 μL of Debris Removal Solution, and 260 μL of PBS was layered on it. After removing supernatant, the pellet was resuspended with 200 μL of PBS including 0.2% BSA and subjected to FACS with the FACSAria instrument (Becton Dickinson). Gating strategies are presented in Extended Data Fig 7a.

### Live imaging in cortical slice culture

An *in vivo* imaging was modified as previously described^17^. Live imaging of cortical slice culture was performed using forebrains isolated from *Cx3cr1-EGFP*^+/-^;*H2B-mCherry*^+/-^ mice at P3. The forebrains were embedded in 2% agarose gel and sliced at 400 nm using a vibratome. These brain slices were placed on the glass-bottom dishes (Iwaki, Cat#3910-035), fixed in collagen gel (Cellmatrix Type I-A [Collagen, Type I, 3 mg/mL, pH 3.0], Nitta Gelatin, Cat#KP-2100), and added in a medium composed of DMEM/F12 (Sigma) containing 5% FBS (Invitrogen), 5% horse serum (HS) (Invitrogen), penicillin/streptomycin (50 U/ml, each) (Meiji Seika Pharma Co., Ltd.), N-2 Supplement (Thermo Fisher Scientific, Cat#17502001), and B-27 Supplement without vitamin A (Thermo Fisher Scientific, Cat#12587010). Time-lapse *in vivo* imaging was conducted using a CSU-X1 confocal microscope (Yokogawa, Tokyo, Japan) with 40x (Olympus LUMPLanFL N 40x / 0.80W) objective. Chambers for culture were filled with 40% O_2_ and 5% CO_2_. The Z-stack images (z-stack pitch;1.5 μm) were acquired using MetaMorph software (Molecular DEVICES). Each image is taken every 5 min. The z-stack images were expanded using a MATLAB program, and the appropriate MNs were identified in each image by analyzing above and below images individually.

### RNA-seq analysis

The library for RNA-seq analysis was constructed with SMART-seq Stranded Kit (Takara) using 10,000 isolated cells by FACS and subjected to sequencing on the Illumina HiseqX platform to yield 151-base paired end reads. Approximately 10 to 20 million sequences were obtained from each sample. Sequences were mapped to the reference mouse genome (mm10) with hisat2 software (10.1038/s41587-019-0201-4). Only uniquely mapped fragments with no base mismatches were used for the analysis. Gene expression was quantitated as fragments per kilobase of exon per million mapped reads (FPKM) on the basis of Genecode M20 (mm10). DEGs between GFP^Neg^, GFP^Low^, and GFP^High^ groups were determined as genes whose FDR were lower than 0.05 with a paired-comparison experiment of *edgeR* of the *R* package. Based on FPKM value, the genes showing GFP^Low^/GFP^Neg^>2, GFP^Low^-GFP^Neg^>50 in GFP^Low^ microglia and GFP^High^/GFP^Neg^>5, GFP^High^-GFP^Neg^>200 in GFP^High^ microglia were defined as upregulated genes following MNs incorporation. The enrichment analyses were conducted using Enrichr analysis tool (https://maayanlab.cloud/Enrichr/).

## Data availability

The sequence data have been deposited in the DNA Data Bank of Japan (DDBJ) Sequence Read Archive under the accession codes: DRA015927.

## Statistical analysis, image processing, and omics data analysis

The statistical data were calculated by using GraphPad Prism (GraphPad Software) compared by two-tailed Student’s *t*-test and one-way ANOVA. Adobe Creative Could CC (Photoshop 22.1.0 and Illustrator 25.0.1) (https://www.adobe.com/), FIJI Image J 2.1.0/1.53c (https://imagej.nih.gov/ij/), and MATLAB (https://www.mathworks.com/) were used for all image processing. The GO analysis obtained by proteomics screening was conducted by Cytoscape v3.9.1.(https://cytoscape.org/) installed with Clue GO v2.5.9. plugin software.

## Acknowledgments

We thank the lab members for helpful discussions and technical supports. We thank Dr Mineko Kengaku (iCeMS, Kyoto University) for technical advice of the neuronal migration experiments. We thank Dr Makoto Urushitani (Shiga University of Medical Science) for distributing BV2 cells. The mass spectrometry analysis was supported by "Nanotechnology Platform Project" operated by the Ministry of Education, Culture, Sports, Science and Technology (MEXT), Japan (No. JPMXP09S19NM0031). We thank Dr. Louis J Irving (Univ. of Tsukuba) for editing of this manuscript. This work was supported by Grant-in-Aid from the Ministry of Education, Science, Sports and Culture of Japan JSPS KAKENHI (16KK0158 (FT), 20K05951, Kao foundation for health science (F.T), Gout and uric acid foundation (F.T), JP20H05688 (KN)), JSPS Research Fellowship for Young Scientists (19J20619 (SY)).

## Author contributions

S.Y. and F.T. designed research; S.Y., N.A., Y.K., H.K., K.A., A.K., K.H., K.K., T.O., B.S., and F.T. performed research; S.Y., N.A., Y.K., H.K., K.A., A.K., M.S.,T.N. and F.T. analyzed data; T.N developed MATLAB program; Y.K., Y.G., and I.T. provide materials. S.Y and F.T. wrote the paper. Study supervision: Y.G., K.N., T.C. and F.T

**Extended Data Fig. 1**

(a) Immunostaining of MAP2^+^ neurons. Primary cultured neurons (5 DIV) were stimulated with the recombinant 100 ng/ml Reelin for 6 hours. Scale bar: 10 μm. (b) The graph shows the percentage of MN^+^ neurons. 10 fields, n=3 independent samples, mean ± SEM, *p* value was calculated by Student’s *t*-test. (c) Schematic illustration for injecting recombinant Reelin. (d) The recombinant Reelin (100 ng) was stimulated into the mice brain for 6 hours, and the brain sections were stained using DAPI. P7, Scale bar: 20 μm. (e) The graph shows the percentage of MNs. 15 images obtained from three independent brain, mean ± SEM, *p* value was calculated by Student’s *t*-test. (f) Schematic illustration for an *in vitro* migration assay. (g) Immunostaining of migrated neurons at the bottom sides of the trans-well. Nuclear shapes were detected by Lamin B1 staining. Asterisks indicate the marks of the hole. Arrow indicates the micronucleus. Scale bar: 10 μm. Illustration indicates the condition of migration, herniation, and micronuclear formation. (h) Immunostaining of migrated neurons at the bottom side of the trans-well. Yellow arrows indicate MNs. Scale bar: 10 μm. (i)(j) The graph shows the population of MN^+^ neurons.(i) n=6, (j) n=3 trans-wells. More than 200 neurons were analyzed from one trans-well. (i) pore size; 3 μm, (j) pore size 12 μm, mean ± SEM, *p* value was calculated by Student’s *t*-test. (k) Schematic illustration for an *in vitro* mechanical stress assay. Neuro2A cells received mechanical stresses by pumping. (l) Neuro2A cells were stained with Hoechst. MNs were generated by mechanical stress. Scale bar: 10 μm (m) The graph shows the percentage of population of MN^+^ cells. Low; 5.0 x 10^4^ cells/300μl, High; 5.0 x 10^4^ cells/150μl, n=9 images, mean ± SEM, *p* values were calculated by one-way ANOVA Dunnett’s multiple comparison test.

**Extended Data Fig. 2**

(a) Immunostaining of MAP2^+^ neurons in the cerebral cortex of either WT or *NexCre:Atg7^f/f^* at P21. Yellow arrows indicate MNs. Scale bar: 20 μm. (b) The graph shows the population of MNs in the cerebral cortex. n=20 images, 10 images obtained from two independent brain; mean ± SEM, *p* value was calculated by Student’s *t*-test. (c) Immunostaining of Iba1^+^ microglia in the cerebral cortex of either WT or *NexCre:Atg7^f/f^* at P21. Yellow arrows indicate MNs. Scale bar: 20 μm. (d) The graph shows the percentage of MNs^+^ microglia in the cerebral cortex. Data were combined from 3 independent brains (8 images per brain, total 24 images), mean ± SEM, *p* value was calculated by Student’s *t*-test. (e) Immunoblotting of the extracellular MNs (P15) obtained from primary cortical neurons in the presence or absence of 100 nM BafA treatment. (f) Primary cortical neurons were stimulated with either 300 nM BafA for 3 hours or 50 μM etoposide (Eto, positive control for inducing apoptosis) for 24 hours. Immunostaining of cleaved-caspase 3 and MAP2^+^ cortical neurons (5 DIV). Scale bar: 50 μm. (g) The graph shows the fluorescence intensity of cleaved-caspase 3 with MAP2. mean ± SEM, *p* values were calculated by one-way ANOVA Dunnett’s multiple comparison test. (h) Schematic illustration of the collection of extracellular micronuclear for immunocytochemistry. (i) Immunostaining of Tubulin and Rab35 resided in the extracellular MNs. Scale bar: 20 μm. (j) Immunostaining of transfected TCP subunits in the migrated neurons at the bottom side of the trans-well (3 DIV). Yellow arrows indicate MNs. Scale bar: 10 μm. (k) Immunostaining of GFP^+^-MNs in the cerebral cortex. A small amount of Tubulin resided in GFP^+^-MNs. P14. Scale bar: 10 μm. (l) Immunostaining of GFP^+^ MNs in the cerebral cortex. MAP2 and LC3 resided in GFP^+^-MNs. P14. Scale bar: 20 μm (upper) and 5 μm (bottom). (m) Working hypothesis of micronuclear secretion dependent on autophagy-lysosome machinery. TCP subunit is a potential receptor linking the micronucleus to LC3.

**Extended Data Fig. 3**

(a) Schematic illustration for an *in vitro* micronuclear transfer assay. (b) TEM image of BV2 cells after treatment with conditioned medium. NE: nuclear envelope, MN: micronucleus, Cyto: cytoplasm. Scale bar: 1.0 μm. (c) Immunostaining of BV2 cells with GFP^-^-and GFP^+^-MNs. Scale bar: 20 μm. (d) Immunostaining of GFP^-^- and GFP^+^-MNs in BV2. Yellow arrow indicates micronucleus. Scale bar: 10 μm. (e) The graph shows the percentage of neuronal MNs (GFP^+^ MNs) containing BV2 cells after treatment with a conditioned medium. Each spot indicates the combined value of 10 images from one experiment. n=3 independent experiments. NB; fresh neurobasal medium, mean ± SEM, *p* values were analyzed by one-way ANOVA Dunnett’s multiple comparisons test. (f) Schematic illustration for eliminating MNs. (g) The graph shows MNs^+^ BV2 cells after treatment with a conditioned medium. mean ± SEM, *p* value was calculated by one-way ANOVA Tukey’s multiple comparisons tests. Each spot indicates the combined value of more than 9 images from one experiment. Each column indicates a set of 3 independent experiments. (h) Live imaging of *Cx3cr1-EGFP*;*H2B-mCherry* cortical slice culture (P3). The interval of taking images was every 5 min. Yellow arrowheads indicate MN in the microglia. Cyan asterisks indicate MN out of the microglia. No.1-6, 89 z-stack images/field, No. 7-24, 92 z-stack images/field, 1.5 µm pitch. Scale bar: 10 μm.

**Extended Data Fig. 4**

(a) Tailing immunostained images of Iba1^+^ microglia in the cerebral cortex. WT, P14. Scale bar: 20 μm. (b) Representative images of microglia in Layer 1 (L1) and Layers 2-6 (L2-6) from the insets of Extended Data Fig. 4a. Scale bar: 20 μm. (c)(d) The graphs show the analyses of microglial characters. 3 brains. (c) Process length, (d) the number of the process. mean ± SEM, *p* value was calculated by Student’s *t*-test.

**Extended Data Fig. 5**

(a) Immunostaining of Iba1^+^ and CD68^+^ microglia with or without MNs (MN^-^; micronuclei-negative, MN^+^; micronuclei-positive). P14, Scale bar: 10 μm. (b) The graph shows the expression level of CD68. The numbers counted are shown in the graph. 5 brains. mean ± SEM, *p* values were analyzed by one-way ANOVA Dunnett’s multiple comparisons test. (c) Immunostaining of Iba1^+^ and Tmem119^+^ microglia with or without MNs. P14, Scale bar: 10 μm. (d) The graph shows the expression level of Tmem119. The numbers counted are shown in the graph. 3 brains. mean ± SEM, *p* values were analyzed by one-way ANOVA Dunnett’s multiple comparisons test. (e) Immunostaining of Iba1^+^ and P2ry12^+^ microglia with or without MNs. The numbers counted are shown in the graph. P14, Scale bar: 10 μm. (f) The graph shows the expression level of P2ry12. 5 brains, mean ± SEM, *p* values were analyzed by one-way ANOVA Dunnett’s multiple comparisons test.

**Extended Data Fig. 6**

(a) The graph shows the expression level of cGAS mRNA in each cell (ASC; astrocyte, NEUR; neuron, OPC; oligodendrocyte precursor cell, iOLG; immature oligodendrocyte, mOLG; myelinating oligodendrocyte, MG/MAC; microglia/macrophage, EC; Endothelial cell). This data was obtained from the Brain RNA-Seq database (https://www.brainrnaseq.org/). (b) Immunostaining of Iba1^+^ microglia in the cerebral cortex of either WT or *cGAS^-/-^* mice at P6. Scale bar: 20 μm. (c)-(e) The graphs show the analyses of microglial state. 2 brains. The numbers in the graph represent the numbers of cells counted. (c) The number of processes, (d) process length, (e) expression level of CD68. mean ± SEM, *p* values were analyzed by Student’s t-test. (f) Immunostaining of CD68 in Iba1^+^ microglia of either WT or *cGAS^-/-^* mice, P6, Scale bar: 10 μm. (g) The graph show the expression level of CD68. The numbers in the graph represent the numbers of cells counted. mean ± SEM, 2 brains. *p* values were analyzed by one-way ANOVA Tukey’s multiple comparisons test. (h) Immunostaining of CD68 in Iba1^+^ microglia of either WT or *cGAS^-/-^*mice, MN^-^; micronuclei-negative, MN^+^; micronuclei-positive. P14, Scale bar: 10 μm. (i) The graph show the expression level of CD68. The numbers in the graph represent the numbers of cells counted. mean ± SEM, 2 brains. *p* values were analyzed by one-way ANOVA Tukey’s multiple comparisons test.

**Extended Data Fig. 7**

(a) Representative gating strategy for microglia in *NexCre:LSL-Sun1GFP* mice. (b) The GO enrichment analysis of DEG in GFP^High^ MN^+^ microglia (1,663 genes). Data was analyzed by Enrichr analysis tool. (c) The scatter plot indicates the ratio and differences of FPKM between GFP^Neg^ MN^-^ microglia and GFP^High^ MN^+^ microglia. FPKM: GFP^High^/GFP^Neg^>5, GFP^High^-GFP^Neg^>200, (d) The Heat map indicates the comparison of FPKM among GFP^Neg^ MN^-^ microglia, GFP^Low^ MN^+^ microglia, and GFP^High^ MN^+^ microglia.. FPKM: GFP^High^/GFP^Neg^>5, GFP^HIgh^-GFP^Neg^>200 (e)-(r) The graph shows the expression of each gene obtained from bulk-RNAseq analysis in GFP^High^ MN^+^ microglia. (ASC; astrocyte, NEUR; neuron, OPC; oligodendrocyte precursor cell, iOLG; immature oligodendrocyte, mOLG; myelinating oligodendrocyte, MG/MAC; microglia/macrophage, EC; Endothelial cell). This data was obtained from the Brain RNA-Seq database (https://www.brainrnaseq.org/).

**Extended Data Fig. 8**

(a)-(k) The graphs show the value of FPKM obtained from bulk-RNAseq analysis in GFP^Low^ MN^+^ microglia. Top 11 genes out of 591 genes are represented, FPKM: GFP^Low^/ GFP^Neg^>2, GFP^Low^-GFP^Neg^>50. (l)-(v) The graph shows the expression of each gene obtained from bulk-RNAseq analysis in GFP^High^ MN^+^ microglia. (ASC; astrocyte, NEUR; neuron, OPC; oligodendrocyte precursor cell, iOLG; immature oligodendrocyte, mOLG; myelinating oligodendrocyte, MG/MAC; microglia/macrophage, EC; Endothelial cell). This data was obtained from the Brain RNA-Seq database (https://www.brainrnaseq.org/).

