## Extended Data Figures for "Propagation of neuronal micronuclei regulates microglial states"

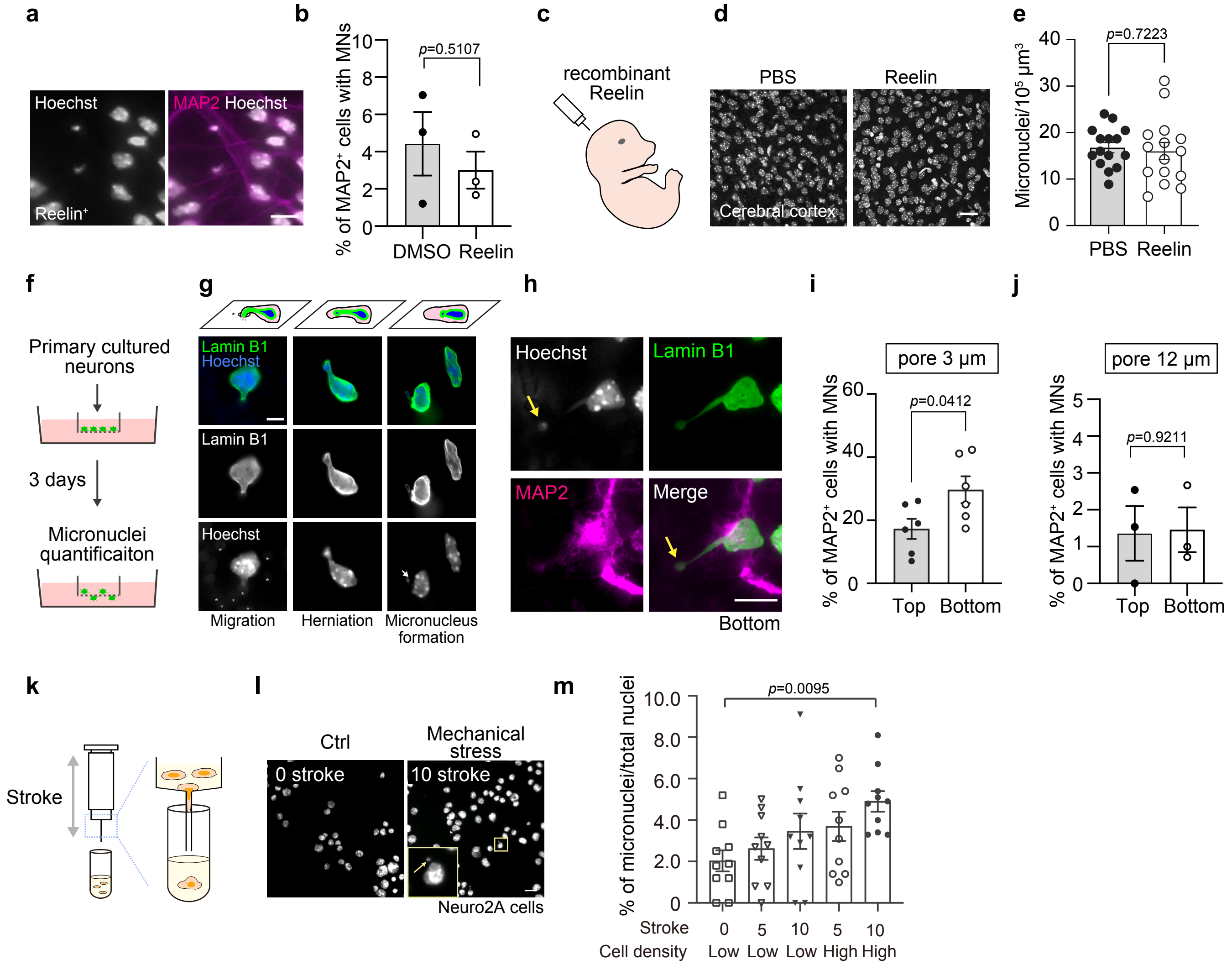


**Extended Data Fig. 1**

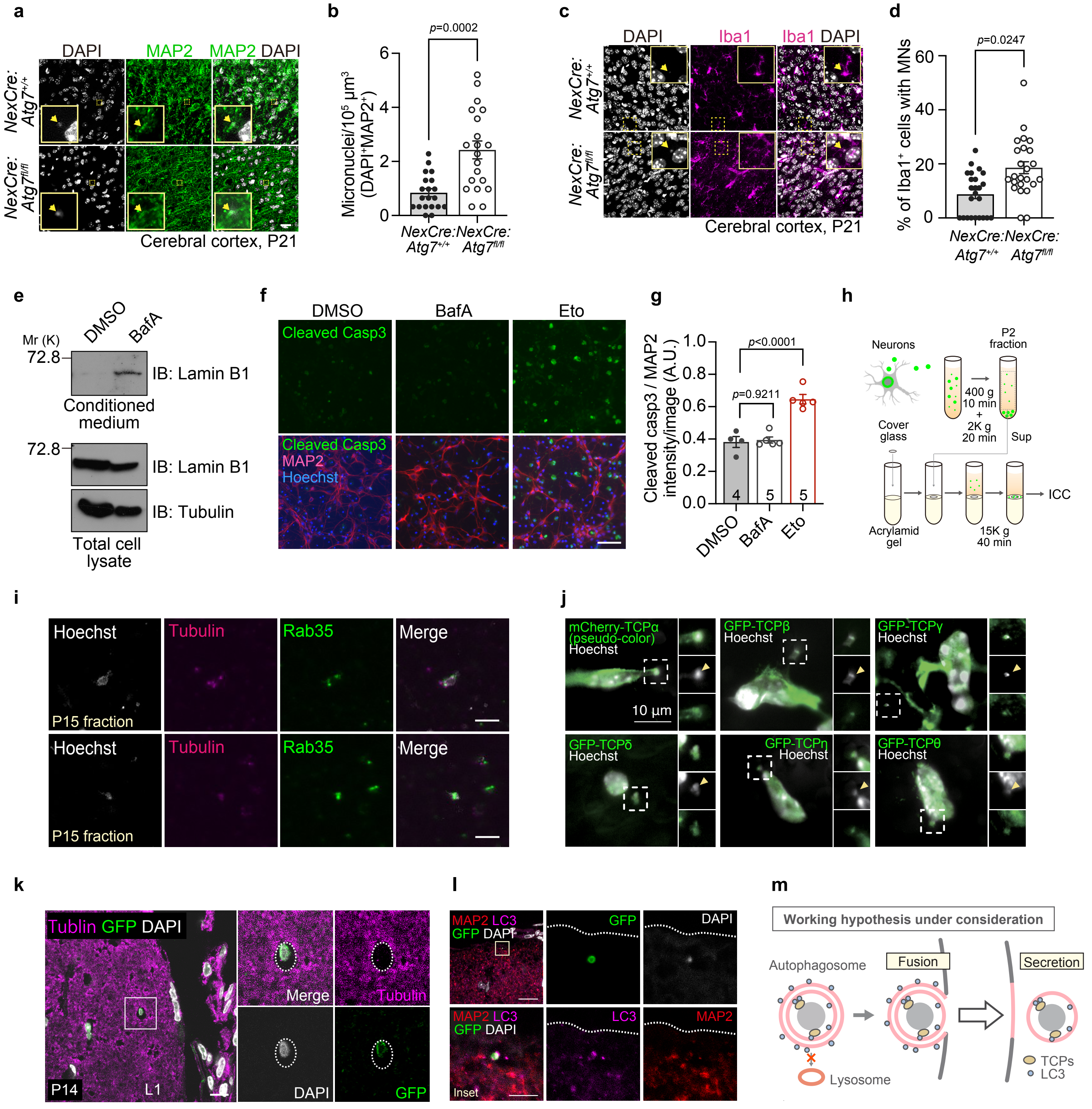


**Extended Data Fig. 2**

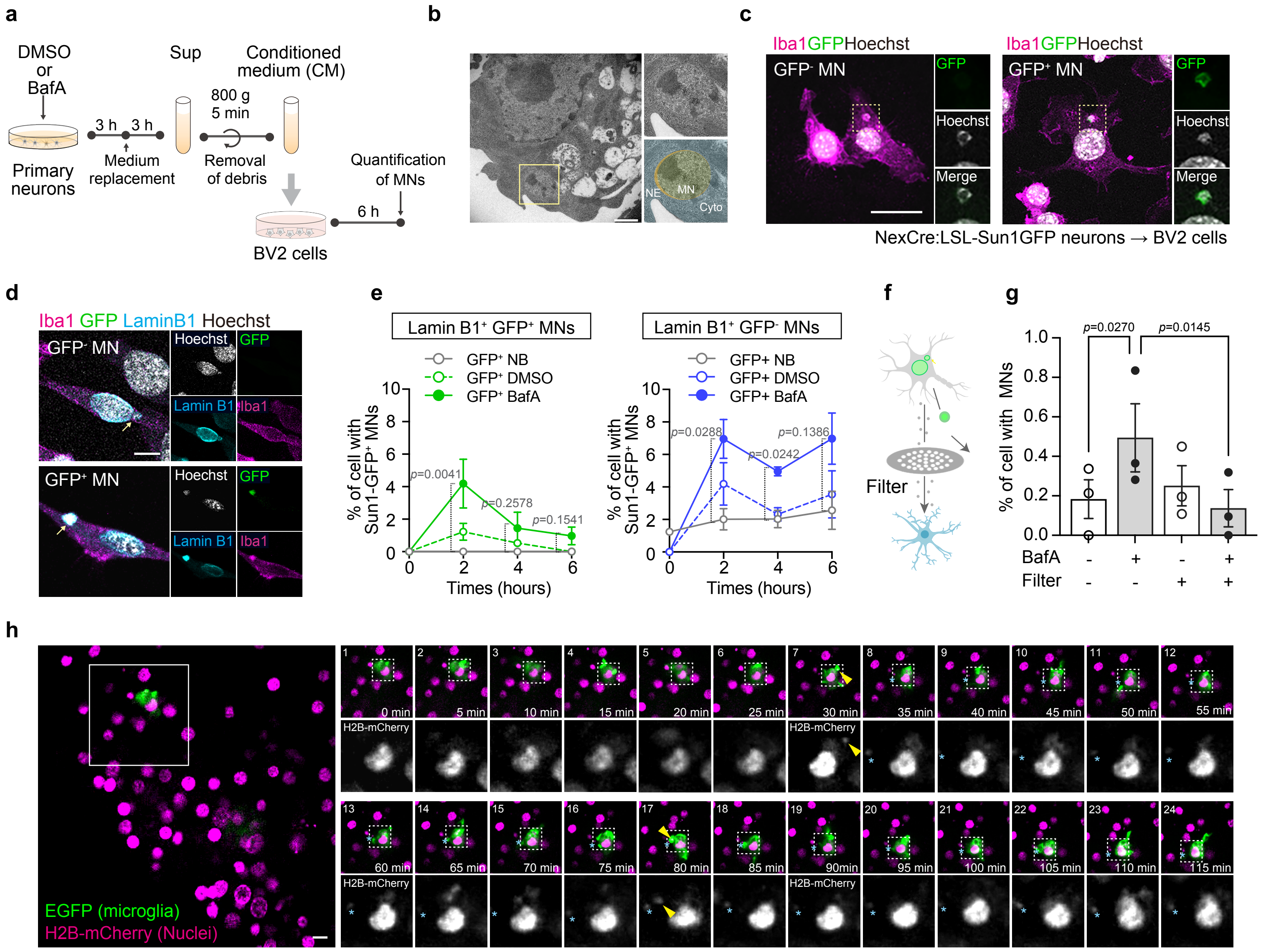


**Extended Data Fig. 3**

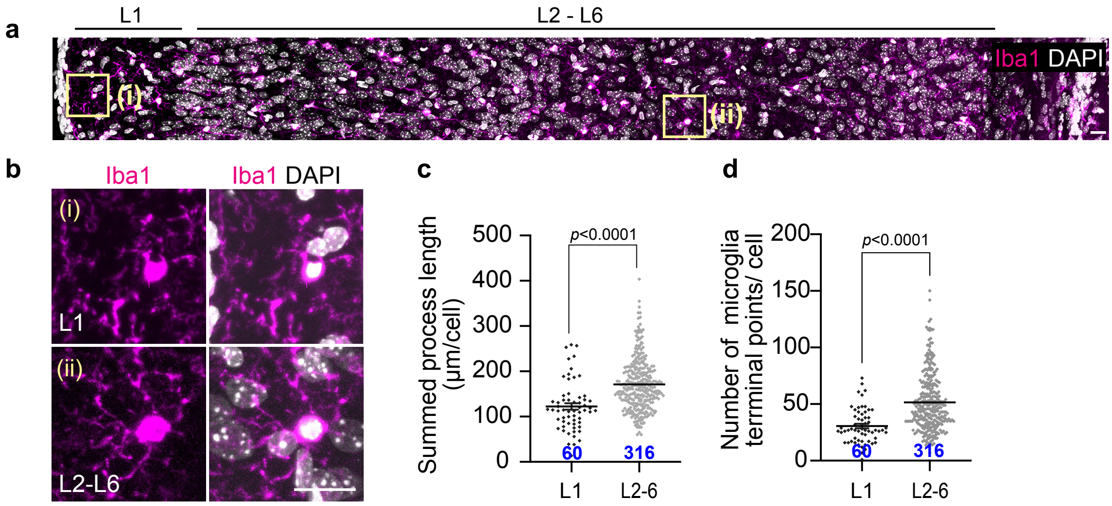


**Extended Data Fig. 4**

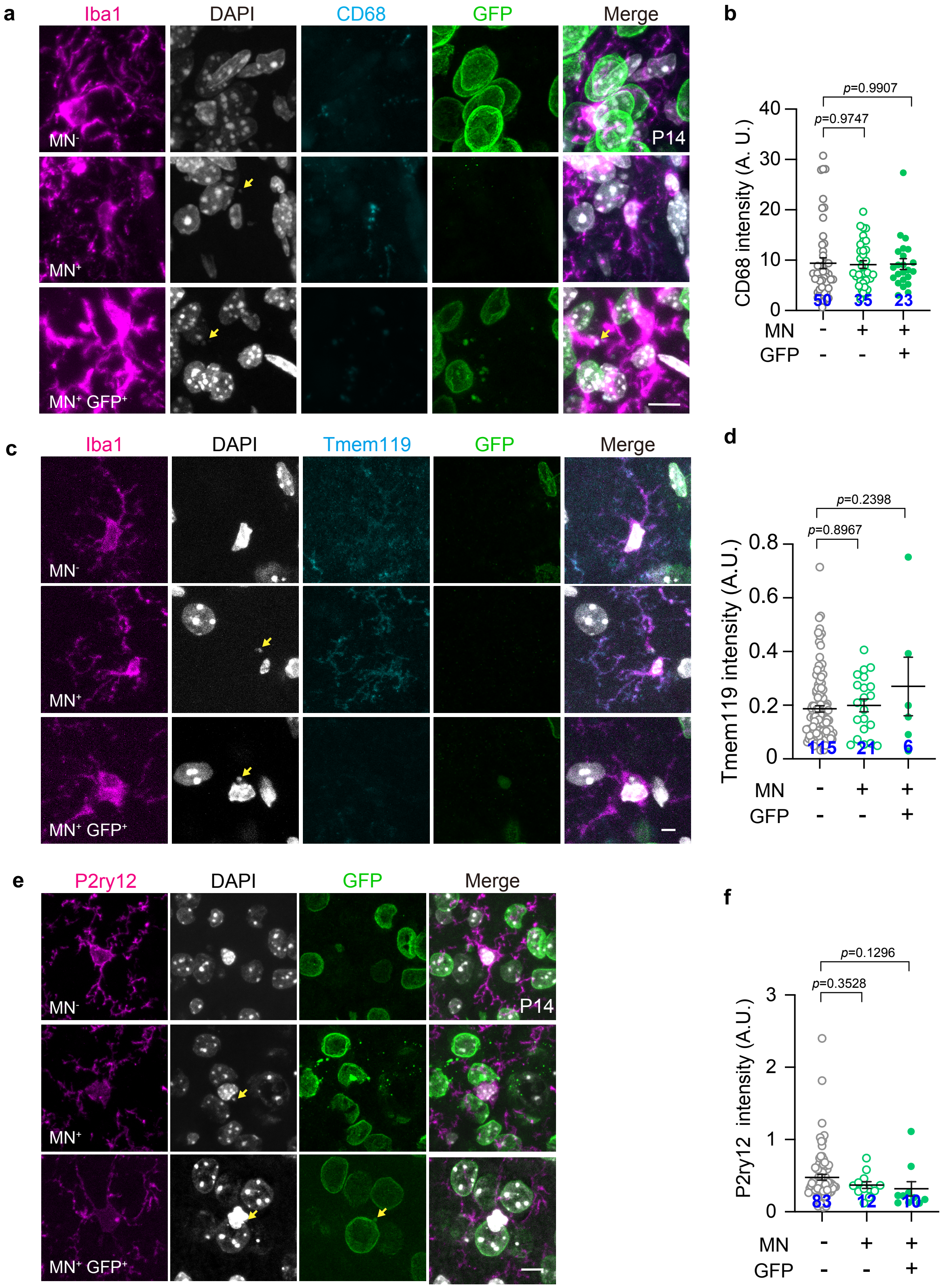


**Extended Data Fig. 5**

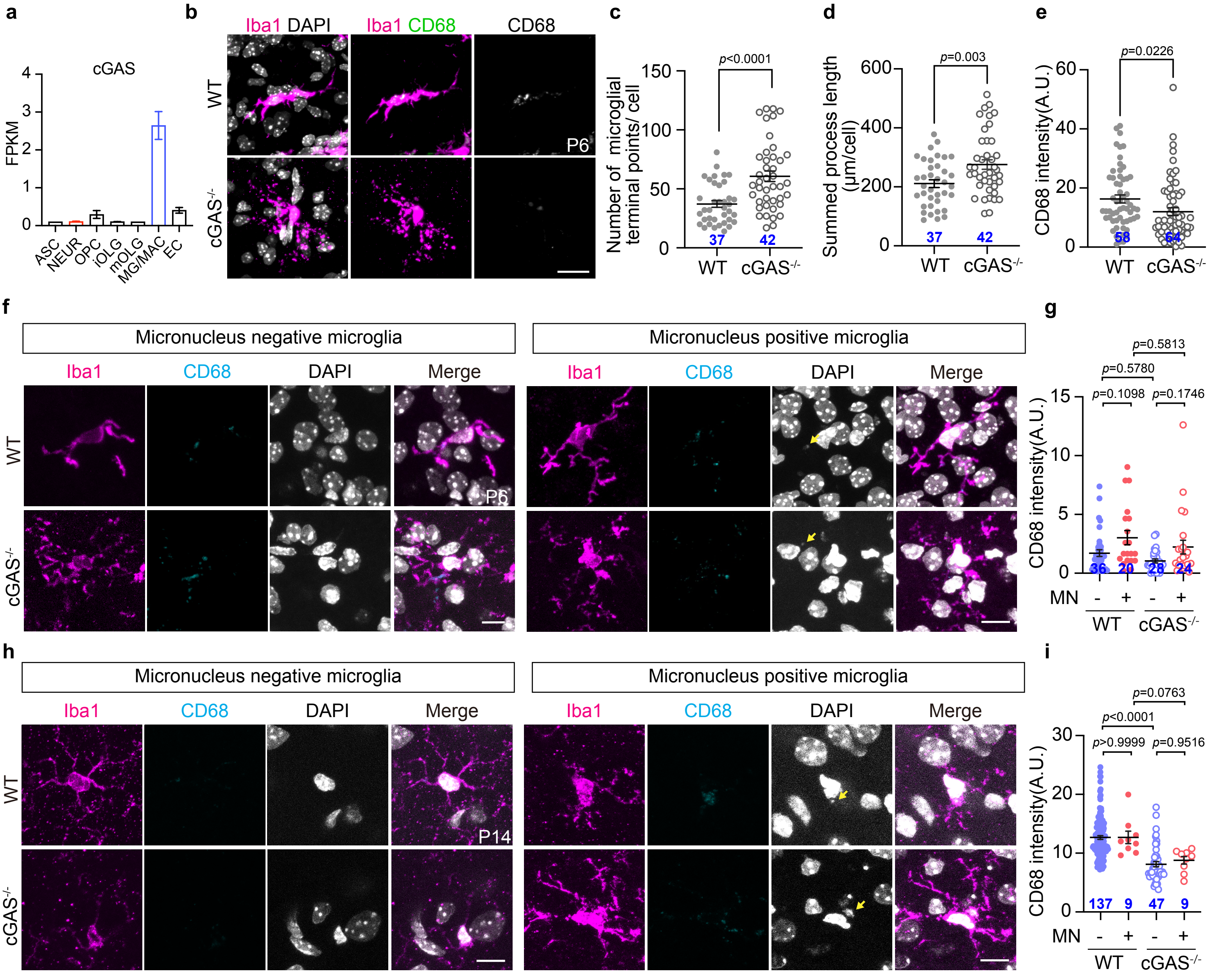


**Extended Data Fig.6**

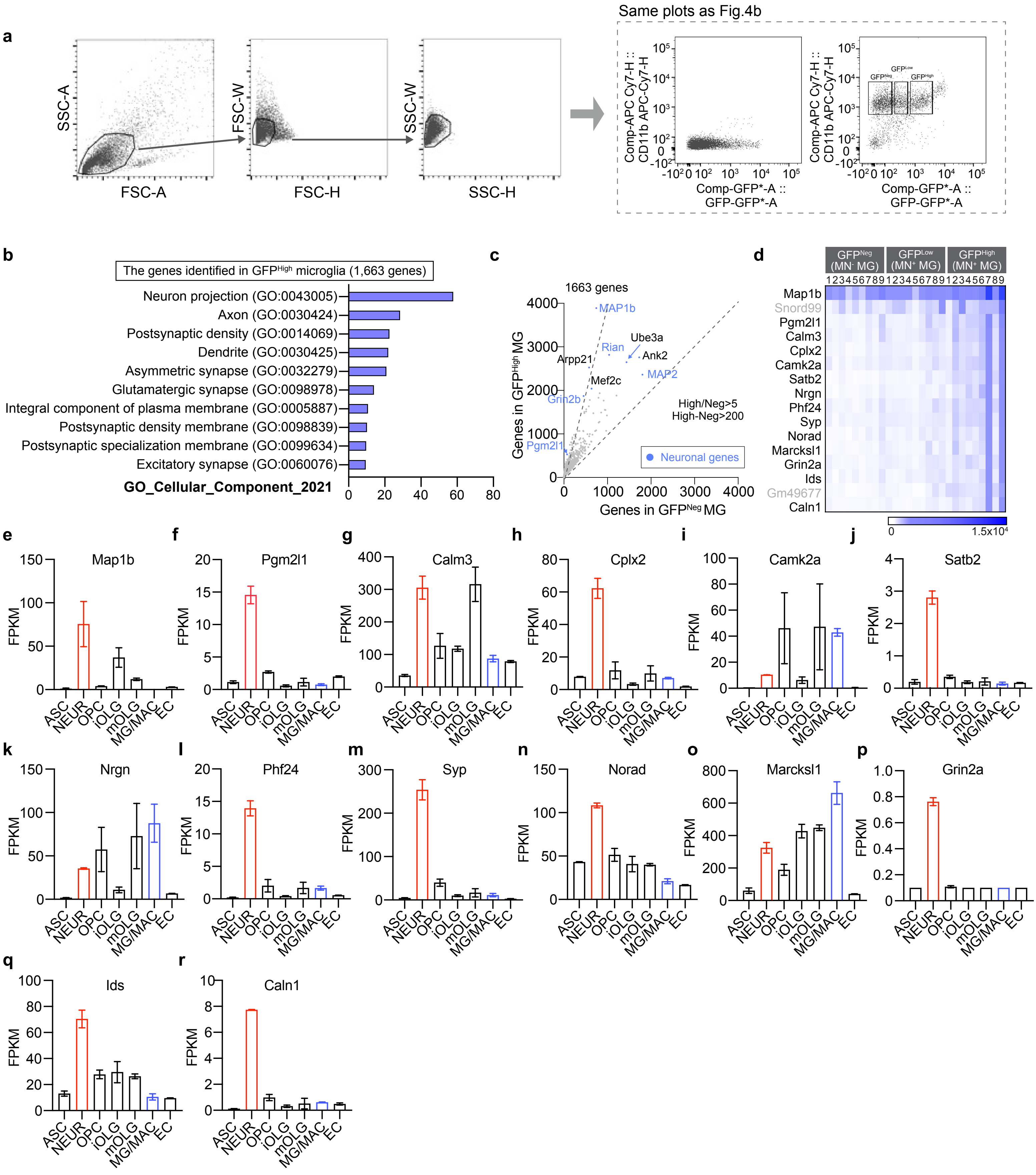


**Extended Data Fig.7**

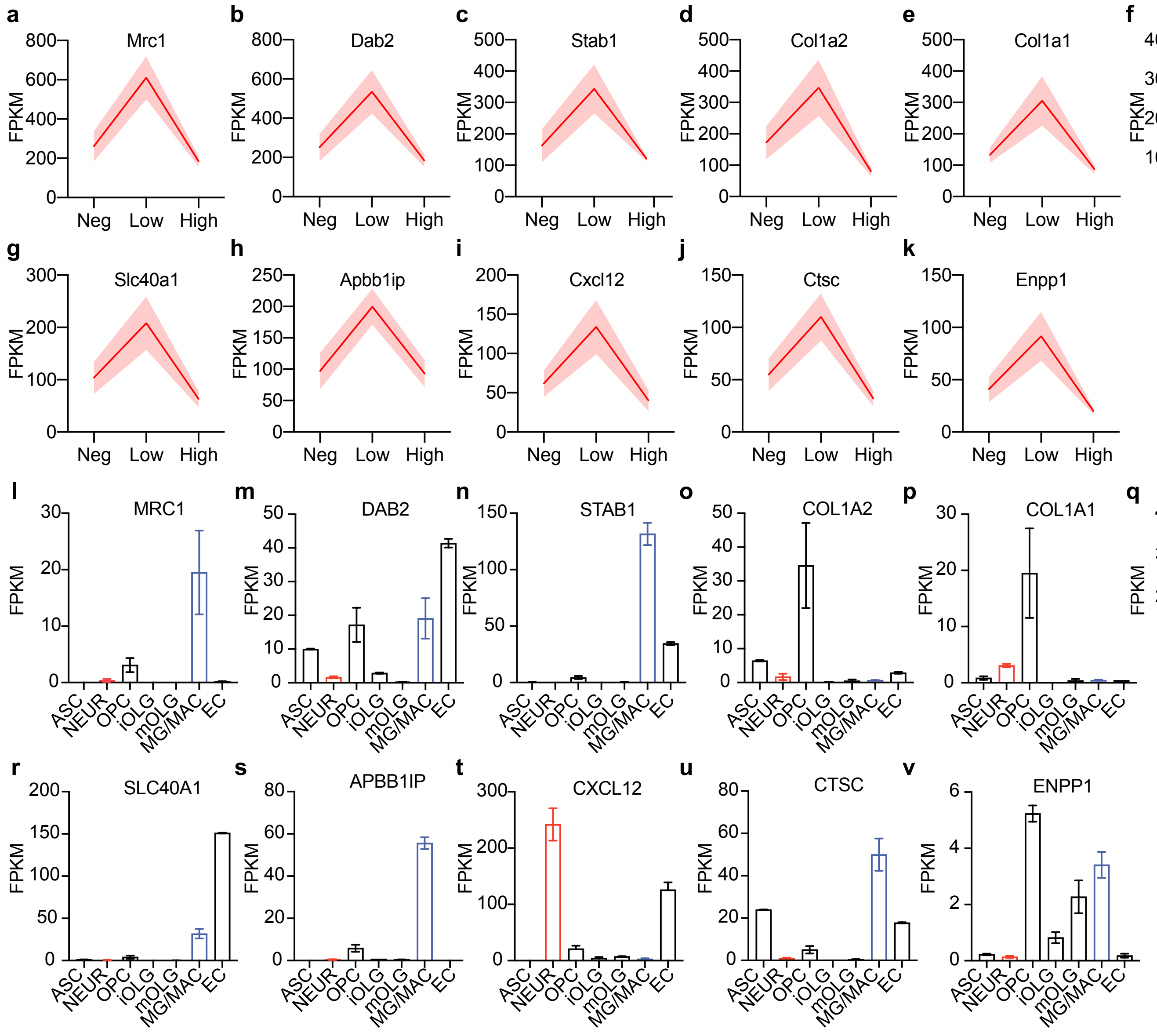


**Extended Data Fig.8**
